# Application of the *in vivo* oxidative stress reporter Hmox1 as mechanistic biomarker of arsenic toxicity

**DOI:** 10.1101/2020.06.19.161117

**Authors:** Francisco Inesta-Vaquera, Panida Navasumrit, Colin J. Henderson, Tanya G. Frangova, Tadashi Honda, Albena T. Dinkova-Kostova, Mathuros Ruchirawat, C. Roland Wolf

## Abstract

Inorganic arsenic (iAs) is a naturally occurring metalloid present in drinking water and polluted air exposing millions of people globally. Epidemiological studies have linked iAs exposure to the development of numerous diseases including cognitive impairment, cardiovascular failure and cancer. Despite intense research, an effective therapy for chronic arsenicosis has yet to be developed. Laboratory studies have been of great benefit in establishing the pathways involved in iAs toxicity and providing insights into its mechanism of action. However, the *in vivo* analysis of arsenic toxicity mechanisms has been difficult by the lack of reliable *in vivo* biomarkers of the effects of iAs. To resolve this issue we have applied the use of our recently developed stress reporter models to study iAs toxicity. The reporter mice Hmox1 (oxidative stress/inflammation; HOTT) and p21 (DNA damage) were exposed to iAs at acute and chronic, environmentally relevant, doses. We observed induction of the oxidative stress reporters in several cell types and tissues, which was largely dependent on the activation of transcription factor NRF2. We propose that our HOTT reporter model can be used as a surrogate biomarker of iAs-induced oxidative stress, and it constitutes a first-in-class platform to develop treatments in arsenicosis. Indeed, in a proof of concept experiment, the HOTT reporter mice were able to predict the therapeutic utility of the antioxidant *N*-acetyl cysteine in the prevention of iAs associated toxicity.

## Introduction

The World Health Organization recognizes that at least 140 million people in 50 countries have been drinking water containing inorganic arsenic (iAs), a class 1 carcinogen, at levels above the recommended guideline value of 10 μg/L [1]. In some countries the maximum allowable level of iAs in drinking water has been reviewed, as different studies have found increased risk of disease development at concentrations below 10 μg/L [2, 3] [4]. Recently, as a direct consequence of increased industrial activity and traffic pollution, the concentrations of iAs in polluted air (e.g. aggregated to particulate matter) has become another significant health risk to exposed populations [5]. Exposure to iAs at different stages of development (e.g. *in utero*, childhood or adulthood) leads to a number of health complications at birth and later in life, such as cardiotoxicity (ischemia, arrhythmia and heart failure) [6], neurotoxicity (cognitive impairment, memory defects) [7] and cancer (skin, bladder and lung) [1]. In human populations, the postulated mechanisms of toxicity of iAs include oxidative stress, DNA damage and inflammation. This is reflected in a variety of biomarkers used to detect iAs exposure. For example, lipid peroxides are significantly higher in highly exposed populations (360 ± 173 μg As/L in their urine) compared with less exposed subjects (71 ± 13 μg As/L) [8]. In low-arsenic (< 10 μg/l) exposed individuals, the levels of urinary 8-nitroguanine and salivary 8-OHdG markers of oxidative stress are increased [9] [10], whereas in high-arsenic exposed individuals (90–160 μg/l) increases in the levels of urinary 8-OHdG [11] have been reported. Moreover, the individual susceptibility to the toxicity and deleterious effects of iAs may be exacerbated by other variables, such as age [7], capacity to metabolise iAs [12] or a specific exposome [13] [14].

*In vitro* and *in vivo* laboratory studies have been instrumental in providing mechanistic insights, not accessible from epidemiological studies, into the consequences of iAs exposure. Some biomarkers of iAs exposure in humans are used in *in vivo* laboratory experiments, such as serum lipid peroxides [15] or DNA methylation changes [16]. In addition, *in vivo* experimental approaches have been used to understand how arsenic exposure can cause human diseases and particularly cancer [17]. As an example, exposure of adult A/J mice (a strain with relatively high spontaneous adenoma/adenocarcinoma incidence) to 1, 10 or 100 ppm iAs for 18 months resulted in increased lung tumour incidence [16]. iAs exposure during gestation (a highly sensitive period for carcinogenesis) has been linked to cancer development later in life [18]. For example, C3H mice exposed to 42.5 or 85 ppm iAs in drinking water for a brief period during gestation produced a dose-dependent induction of tumours in the liver, adrenal, lung and ovary when the offspring reached adulthood [19]. In CD1 mice, whole-life iAs exposure (from preconception, through embryo development, lactation, and up to 104 weeks old) also resulted in a higher tumour incidence [20].

Despite intense investigations, the translation of laboratory findings into human disease outcomes still requires research [21]. It is increasingly recognized that both experimental and biological factors (e.g. mouse genetic backgrounds, dose/timing of iAs exposure, sex or coexposure to other environmental components) are determinants of the *in vivo* toxicity of iAs. One limitation in the study of iAs toxicity *in vivo* is the absence of reliable biomarkers, with the majority of the *in vivo* studies relying on generic measurements of toxicity (such as overall body or tissue weight, lethality), or the products of oxidative stress (malondialdehyde, hydroxynonenal or 8-hydroxyguanosine). Such studies do not provide information about effects on specific tissues, cell types or toxicity pathways. As a consequence, studies of the effects of iAs exposure on embryo development, cancer and pulmonary injury can often report apparently contradictory results [22–28]. Therefore, to advance our understanding of the diseases associated with iAs exposure there is a need to develop new biomarkers and robust animal models targeted at understanding the underlying molecular mechanisms of iAs toxicity [29].

Oxidative stress has been proposed as one of the main mechanisms for iAs toxicity in rodents and humans [30]. The Kelch-like ECH-associated protein 1/nuclear factor erythroid 2 p45-related factor 2 (KEAP1/NRF2) pathway is a major cytoprotective pathway, which is upregulated by iAs *in vivo* [31, 32]. This pathway controls the intracellular redox environment by regulating the expression of about 250 genes involved in antioxidant response, iron catabolism, inflammation and endo and xenobiotic biotransformation. NRF2-KO mice exposed to 100ppm iAs for 6 weeks were more sensitive to arsenic-induced oxidative DNA damage, epigenetic changes and pathological alterations than their wild type counterparts [31]. As a result, the induction of the NRF2 pathway has been proposed as a target to counteract damaging effects of arsenic [31, 33].

We have recently generated a set of next-generation mouse reporter models (the Hmox1 reporter of oxidative stress/inflammation, HOTT) and the p21 reporter (DNA damage), which provide easily measurable readouts of *in vivo* cellular responses to toxic insult, with high-resolution, and in a tissue- and cell-specific manner [34, 35]. In each model, a short viral DNA sequence, known as a 2A sequence [36], is exploited to provide multiple reporters to be expressed (separately) from the endogenous gene promoter. When HOTT reporter mice were exposed to oxidative stress inducers (e.g. paracetamol) or inflammatory mediators (e.g. LPS), a tissue- and cell-specific induction of ß-galactosidase activity was detected [34]. Similarly, when the p21 reporter mice were exposed to DNA damage inducing agents (e.g. γ-irradiation or cisplatin) an increase of p21 expression was detected at a high level of fidelity and resolution in target tissues [35]. These models therefore potentially represent a new approach to understand iAs associated toxicity and to evaluate the therapeutic potential of novel treatments. In this work, we demonstrate that the HOTT reporter models are a robust *in vivo* platform for the detection of iAs associated toxicity in a cell- and tissue-specific manner. In addition, a new reporter line, NRF2-KO_HOTT is described which allows the interrogation of the role of the transcription factor NRF2 in iAs toxicity. The utility of these reporter systems as a biomarker for iAs toxicity *in vivo* was demonstrated by predicting the prevention of iAs toxicity by the antioxidant *N*-acetyl cysteine.

## Materials and Methods

### Animals

All animals used in this study were bred and maintained in the Medical School Resource Unit, University of Dundee, on a C57BL/6N background. We previously described the generation of Hmox-1 (HOTT, HO-1 triple transgenic) and p21 reporter mice [34, 35]. NRF2-KO animals were kindly provided by Prof. John D. Hayes (School of Medicine, University of Dundee) and they have been described before [37]. NRF2-KO_HOTT reporter mice were generated by crossing NRF2-KO animals into HOTT reporter mice. Mice were housed in open-top cages in temperature-controlled rooms at 21°C, with 45-65% relative humidity and 12h/12h light/dark cycle. Mice had *ad libitum* access to food (R&M No.1 for stock females; R&M No. 3 for mating females; arsenic concentrations undetectable or < 1mg/kg, respectively; Special Diet Services, Essex, UK); and water (iAs concentration in water is estimated below the 10μg/l limit). Animals were regularly subjected to health and welfare monitoring as standard (twice-daily). All cages had sawdust substrate and sizzle-nest material provided. Environmental enrichment was provided for all animals.

All animal work described was approved by the Welfare and Ethical use of Animals Committee of the University of Dundee. Those carrying out this work did so with Personal and Project Licences granted by the UK Home Office under the Animals (Scientific Procedures) Act 1986, as amended by EU Directive 2010/63/EU. Animals in study plans were inspected regularly by staff trained and experienced in small animal husbandry, with 24-hour access to veterinary advice.

Animal numbers were guided by power calculations (G*Power; www.gpower.hhu.de), pilot experiments, and previous experience, and experimental design was undertaken in line the 3Rs principles of replacement, reduction, and refinement (www.nc3rs.org.uk).

### Study design

Data in this paper were obtained using female mice, unless otherwise stated. In preliminary experiments, males displayed a comparable activation of the HOTT reporter in an acute exposure model to iAs. Mice were littermates or age-matched to within 3 weeks of each other and ages specified. All animals used in this study were heterozygous for the HOTT or p21 reporter allele unless otherwise specified (HOTT^+/r^ or p21^+/r^). Animals were randomly assigned to control or treatment groups; analysts were not blinded to the identity of biological samples. At the end of studies, all animals were sacrificed by exposure to a rising concentration of CO_2_ and death confirmed by exsanguination.

### Treatments

Sodium arsenite (Sigma, S7400) was dissolved in drinking water and administered by oral gavage for short and transplacental exposure experiments at the indicated final concentrations. For chronic exposure studies, arsenic was administered in the drinking water at concentrations reflecting environmental exposure. The average water intake per animal and per day was 3.7 ml (+/− 0.49 ml) for the 50ppm iAs treatment, and 4 ml (+/- 0.37 ml) for the 5ppm iAs treatment. Water was changed twice per week to avoid arsenic oxidation. All animals appeared healthy throughout the experiment and did not show clinical signs of illness or other potential treatment-related symptoms. Total and organ weights were equivalent between control and exposed animals at necropsy. *N*-acetyl cysteine (NAC; SIGMA) was administered at a concentration of 300mg/kg body weight in water or in combination with iAs by oral gavage (p.o.).

TBE-31, a highly potent NRF2 activator [38], was synthesized as described before [39]. Animals were administered a daily dose of vehicle (corn oil, 5μl/g body weight p.o.) or TBE-31 (5 nmol/g body weight p.o.) for three consecutive days and 24h before tissue harvesting.

### Genotyping

Ear biopsies obtained from mice (4–8 weeks old) were incubated at 55°C overnight in lysis buffer containing 10mM Tris (pH = 8), 75 mM NaCl, 25 mM EDTA, 1% (w/v) SDS and 100 μg ml-1 (39 U mg-1) proteinase K (Sigma). The concentration of NaCl in the reaction was raised to 0.6 M and a chloroform extraction was performed. Two volumes of isopropyl alcohol were added to the extracted supernatant to precipitate genomic DNA (gDNA). A 50 μl volume of TE buffer (10 mM Tris, 1 mM EDTA, pH 8.0) was added to the pellet, and subsequently gDNA was dissolved for 1h at room temperature and stored at 4°C until further use. The typical PCR reaction consisted of a 25 μl volume containing 1.25U Taq DNA polymerase (Thermo-Scientific), 10mM dNTPs, Thermo Taq Buffer, 25 mM MgCl2 and 10 pmol of the primers (HOTT reporter: HO1-KI Fwd, 5_-GCTGTATTACCTTTGGAGCAGG-3_; HO-1-KI Rvr, 5’-CCAAAGAGGTAGCTAATTCTATCAGG-3’); (p21 reporter: p21-KI Fwd, 5’-GCTACTTGTGCTGTTTGCACC-3’; p21-KI Rvr, 5’-TCAAGGCTTTAGGTTCAAGTACC-3’); (Nrf2: Nrf2 Fwd, 5’-TGGACGGGACTATTGAAGGCTG-3’; Nrf2 As Rvr, 5’-GCCGCCTTTTCAGTAGATGGAGG-3’; Nrf2 LacZ Rvr, 5’-GCGGATTGACCGTAATGGGATAGG-3’). The following PCR conditions were applied: 5 min, 95°C initial denaturation; 30 s, 95°C cyclic denaturation; 30 s, 60°C cyclic annealing; 1 min, 72°C cyclic elongation for a total of 35 cycles, followed by a 10 min 72°C elongation step. PCR amplification products were analysed by agarose gel electrophoresis.

### *In vivo* luciferase imaging

Imaging was performed 24h after vehicle or chemical dosing. Reporter mice were injected intraperitoneally (i.p.) with 5 μl g^-1^ body weight RediJect d-Luciferin (PerkinElmer) and anaesthetized by isofluorane before being transferred into the IVIS Lumina II imaging chamber (Perkin-Elmer) for bioluminescence imaging. Luminescence images (automatic exposure, f/stop 1.2, binning 4) and greyscale images (automatic exposure, f/stop 1, binning 2) were acquired. Photon fluxes in regions of interest (ROIs) were quantified using the LivingImage (R) Software, version 4.3.1 (Perkin-Elmer). Luminescence images were rendered using the fire look-up table in ImageJ, and superimposed on greyscale photographs.

### Tissue harvesting and processing for cryo-sectioning

Tissues were rapidly harvested *post mortem* and processed by immersion fixation in 4% *para-* formaldeyde (PFA) (brain, small intestine, skin) for 2h, 3% neutral-buffered formalin (NBF) (liver) for 3h or Mirsky’s fixative (rest of tissues) for 24h. For cryosectioning, tissues were cryoprotected for 24h in 30% (w/v) sucrose in phosphate-buffered saline (PBS) at 4°C. Organs were embedded in Shandon M-1 Embedding Matrix in a dry ice-isopentane bath. Sectioning was performed on an OFT5000 cryostat (Bright Instrument Co.). With the exception of lung (14μm) and brain (20μm) sections, all sections were cut at 10μm thickness. Sections were kept at −20°C until further use. Embryos were dissected in ice-cold PBS and fixed in ice-cold 4% PFA for 2h at 4°C, washed and maintained in PBS before β-galactosidase staining.

### *In situ* β-galactosidase staining

Sections were rehydrated in PBS at room temperature for 15 minutes before being incubated overnight at 37°C in X-gal staining solution: PBS (pH 7.4) containing 2 mM MgCl2, 0.01% (w/v) sodium deoxycholate, 0.02% (v/v) Igepal-CA630, 5 mM potassium ferricyanide, 5 mM potassium ferrocyanide and 1 mg/ml 5-bromo-4-chloro-3-indolyl β-D-galactopyranoside. On the following day, slides were washed in phosphate buffer solution, counterstained in Nuclear FastRed (Vector Laboratories) for 4 min, washed twice in distilled water for 2 minutes and dehydrated through 70% and 95% ethanol (4.5 and 1 minute respectively) before being incubated in Histoclear (VWR) for 3 minutes, air-dried and mounted in DPX mountant (Sigma).

### Hematoxylin – Eosin staining

Sections were deparaffinized in xylene and rehydrated through a graded series of 100–50% (v/v) ethanol solutions, followed by immersion in distilled water. Sections were incubated for 5 min at room temperature in hematoxylin followed by 10 min washing in running tap water. After a 2 min 80% ethanol incubation, samples were incubated in eosin for 10-20 secs. Finally, sections were dehydrated through 95% (v/v) ethanol and two changes of 100% (v/v) ethanol, cleared with two changes of xylene, air-dried, and mounted in DPX medium (Sigma).

### Masson’s staining

Sections were deparaffinized in xylene and rehydrated through a graded series of 100–50% (v/v) ethanol solutions, followed by immersion in distilled water. Sections were sequentially incubated in Bouin’s solution (1h, 56 °C), Celestine blue solution (0.5% w/v Celestine blue, 5% w/v Ferric ammonium sulphate, 12% v/v glycerol; 5 min.), Harris Haematoxylin (5 min), acid alcohol (1% HCl in 70% ethanol; 3 min.), Scott’s tap water (1.4% w/v magnesium sulphate, 2% sodium bicarbonate in distilled water; 5 min), Solution A (0.5% w/v acid fuschin, 0.5% w/v xylidine ponceau in 1% v/v acetic acid; 10 min.), Solution B (1% w/v phosphomolybdic acid; 10 min.), Solution C (2% w/v Light Green SF Yellowish in 2% v/v acetic acid; 10 min). Samples were washed with distilled water for 2 min between incubations. Finally, sections were dehydrated through 95% (v/v) ethanol and two changes of 100% (v/v) ethanol, cleared with two changes of xylene, air-dried, and mounted in DPX medium (all reagents from Sigma).

### Clinical chemistry

Terminal bleeds were collected in heparinized blood collection tubes (Sarstedt). Clinical chemistry assays were performed blind at the clinical pathology laboratory, MRC Harwell.

### Densitometric evaluation

The software ImageJ v1.51q was used for quantitative image analysis of the β-gal staining. The densitometric analysis was based on microscope images that were acquired with a Zeiss Axio Observer (Carl Zeiss, Jena, Germany). A threshold for the red, green and blue (RGB) channels was defined in order to filter the area of blue coloration within the image. The value of each RGB channel of each pixel had to be below the threshold to remove the background in each image and to quantify the corresponding pixels as X-Gal positive. After the software automatically determined the area of blue coloration within the microscopic image, the total area of the image was set in relation to the area of the measured blue staining. Subsequently, the percentage of the area of X-Gal positive staining of each acquired microscope image was determined. Individual values are given for each image quantified.

### Immunoblots

Whole-tissue protein extracts were prepared from snap frozen organs. Briefly, 300 μl of lysis buffer (25mM Tris-HCl pH 7.4, 150mM NaCl, 5mM EDTA, 1% Nonidet P40, 0.5% sodium deoxycholate, 0.1% SDS) supplemented with protease inhibitors and phosphatase inhibitors) was added per 100mg of tissue and homogenized using a Polytron PT2100 benchtop homogenizer rotor. After 30 min on ice, the resulting lysate was centrifuged for 15 min at 13200 rpm on an Eppendorf tabletop centrifuge at 4°C. Supernatant was recovered and protein concentrations were measured by Bradford assay kit (Biorad). SDS-PAGE and immunoblotting was carried out as previously described [40]. Antibodies used included hmox1 (Abcam, ab13243), β-gal (Promega, Z3781), p21 (BD Pharmingen, 556431), NQO1 (Ab2346), GAPDH (Cell signaling, 2118).

### DNA methylation analysis

DNA methylation was analysed by quantitative bisulfite Pyrosequencing assay. Briefly, DNA was isolated from heart and liver tissues according to the manufacturer’s protocol (DNeasy® Blood & Tissue kit, Qiagen). The concentration and purity of total DNA were analyzed using a Nanodrop, ND-1000 spectrophotometer (NanoDrop Technologies, Inc., USA). Genomic DNA (1 μg) was treated with bisulfite using an Epitect Bisulfite kit (Qiagen, Germany), following the manufacturer’s instructions. The bisulfite converted DNA was subjected to quantification of CpG methylation using Pyrosequencing. Two set of primers for gene-specific mouse CpG sites, *DUSP1, EGR1, JUNB* and *SOCS3*, were selected by predesigned PyroMark CpG Assay (Qiagen) *(DUSP1*, Cat. No. PM00294917 and PM00294924; *EGR1*, Cat. No. PM00000525 and PM00305039; *JUNB*, Cat. No. PM00413553 and PM0041360; and *SOCS3*, Cat. No. PM00245245 and PM00245252). PCR reaction mixture contained 0.2 μM of PCR primers, 1x PyroMark PCR Master Mix, 1x CoralLoad Concentrate, 20 ng of bisulfite-converted DNA and RNase-free water in a total volume of 25 μl. The PCR program was performed using an Eppendorf Mastercycler® pro PCR system (Eppendorf, Germany). Subsequently, PCR product (20 μl) was used to perform Pyrosequencing reaction following Pyromark Q96 ID protocols with PyroMark Q96 ID Software 2.5 (version 2.5.10.7). The percentage of methylation at individual CpG sites was quantified by Pyromark CpG software version 1.0.11 (Qiagen, Germany). Samples with a mean methylation above 5% for each CpG site were used for analysis. The levels of methylation in all CpG sites of the *DUSP1, EGR1, JUNB* and *SOCS3* were determined.

## Results

### Effect of arsenic exposure on HO-1 and p21 reporter expression

To interrogate the capacity of the reporter mice to detect iAs-induced stress responses *in vivo*, HOTT reporter mice (HOTT^+/r^) were exposed to iAs orally at low (5.4 μmol/kg) or high (77 μmol/kg) doses. These concentrations are equivalent to an exposure of 6 ppm or 85.71 ppm iAs in drinking water, based on an average water consumption for mice of 3.5 ml/day. The selected doses was chosen to reflect real life environmental exposure (low dose) [41] and also for comparison with literature reports where acute exposure (high dose) was used [17, 42]. Twenty-four hours after the single exposure, a panel of tissues were harvested for histochemical and biochemical analysis (Figure 1 and Figure S1). Basal β-galactosidase activity staining was observed in the brain (hippocampus and cerebellum), spleen (red pulp), lungs (bronchiole, respiratory epithelium) and skin (Figure S5A) as previously described [34]. The basal activity in these tissues was unchanged on iAs exposure. β-galactosidase activity was not detected in the small, large intestine or uterus of exposed mice at any dose. Interestingly, β-galactosidase activity was markedly induced in liver, heart and kidney tissues of mice exposed to 77 μmol/kg arsenic (Figure 1). Hepatocytes in livers and cardiac fibres in heart of exposed mice were consistently positive for LacZ staining in a widespread distribution pattern. In kidneys, the expression was predominantly localized in tubular cells of the cortex and the outer medulla (both inner and outer stripe). Importantly, in mice exposed to the low arsenic dose, a sparse but consistent activation of the reporter in cardiac fibres was observed. To confirm that the iAs-induced reporter activation reflected the expression of the endogenous HO-1 protein, liver and kidney tissues were analysed by immunoblotting (Figure 1B and Figure 1C, respectively). In both tissues, an increase in HO-1 protein levels was observed in mice treated with the high iAs dose. The protein levels of NQO1, an additional NRF2 target gene, were not consistently elevated in tissue lysates of exposed animals. These data are consistent with previous publications and indicate that HOTT mice provide a robust reporter model to detect the consequences of iAs exposure *in vivo* [43]. Specifically, liver, kidney and heart (which are targets for iAs toxicity [12]) were the tissues with high reporter response to iAs exposure.

**Figure 1.**
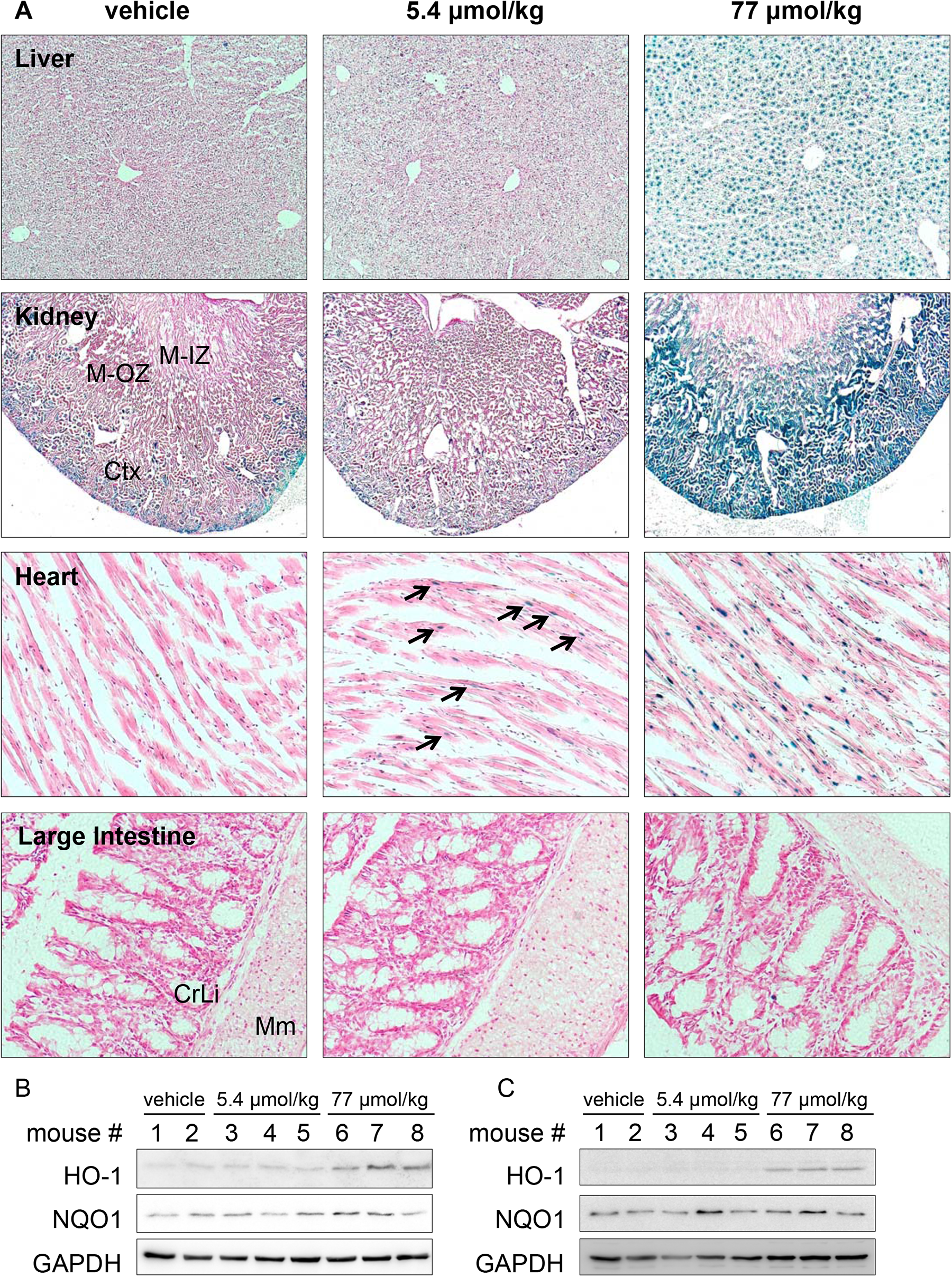
Tissue specific susceptibility to iAs exposure *in vivo* using HOTT reporter mice. Triplicate HOTT^+/r^ mice (female, 16wo aged matched) were dosed (p.o.) with indicated amounts of sodium arsenite. 24h later tissues were harvested and *in situ* β-galactosidase assay performed. **A**. Representative images are shown for liver (10x), kidney (2.5x), Heart (20x) and Large Intestine (10x). Black arrows indicate β-galactosidase activity. Ctx, cortex; M-OZ, medulla outer zone; M-IZ, medulla inner zone; CrLi, crypts of Lieberkuhn: Mm, muscularis mucosa. **B** and **C**. Organ lysates were probed for the indicated proteins. GAPDH was used as a loading control.

Epidemiological studies have linked arsenic exposure *in utero* and in adults to oxidative DNA damage [10, 44]. This can result in toxicity by apoptosis mediated by the p53 pathway [45–47]. We investigated the ability of a high dose of iAs to activate the expression of the p53 target gene p21. In agreement with our previous work, constitutive p21 activation was observed in a number of mouse tissues, including: liver, kidney and heart [35]. A subset of cells types, such as smooth muscle cells in the large intestine and the pulmonary veins, bronchiolar cells in lungs and peritubular myoid cells and spermatids in testis also had constitutive reporter activity (Figure 2 and Figure S2A). Exposure of p21 reporter mice to iAs resulted in no further activation of the reporter in any of the tissues examined. In contrast, western blot analysis of liver and kidney tissue lysates from this experiment demonstrated an induction of HO-1 expression on arsenic treatment, consistent with data obtained in the HOTT mice (Figure 2B and 2C). Also, as a positive control, two male mice carrying the HOTT reporter were exposed to iAs and histochemical analysis showed the expected increase in β-galactosidase activity in liver, kidney and heart tissues (Figure S2B). These results indicate that the initial oxidative stress induced by iAs does not trigger a DNA damage response.

**Figure 2.**
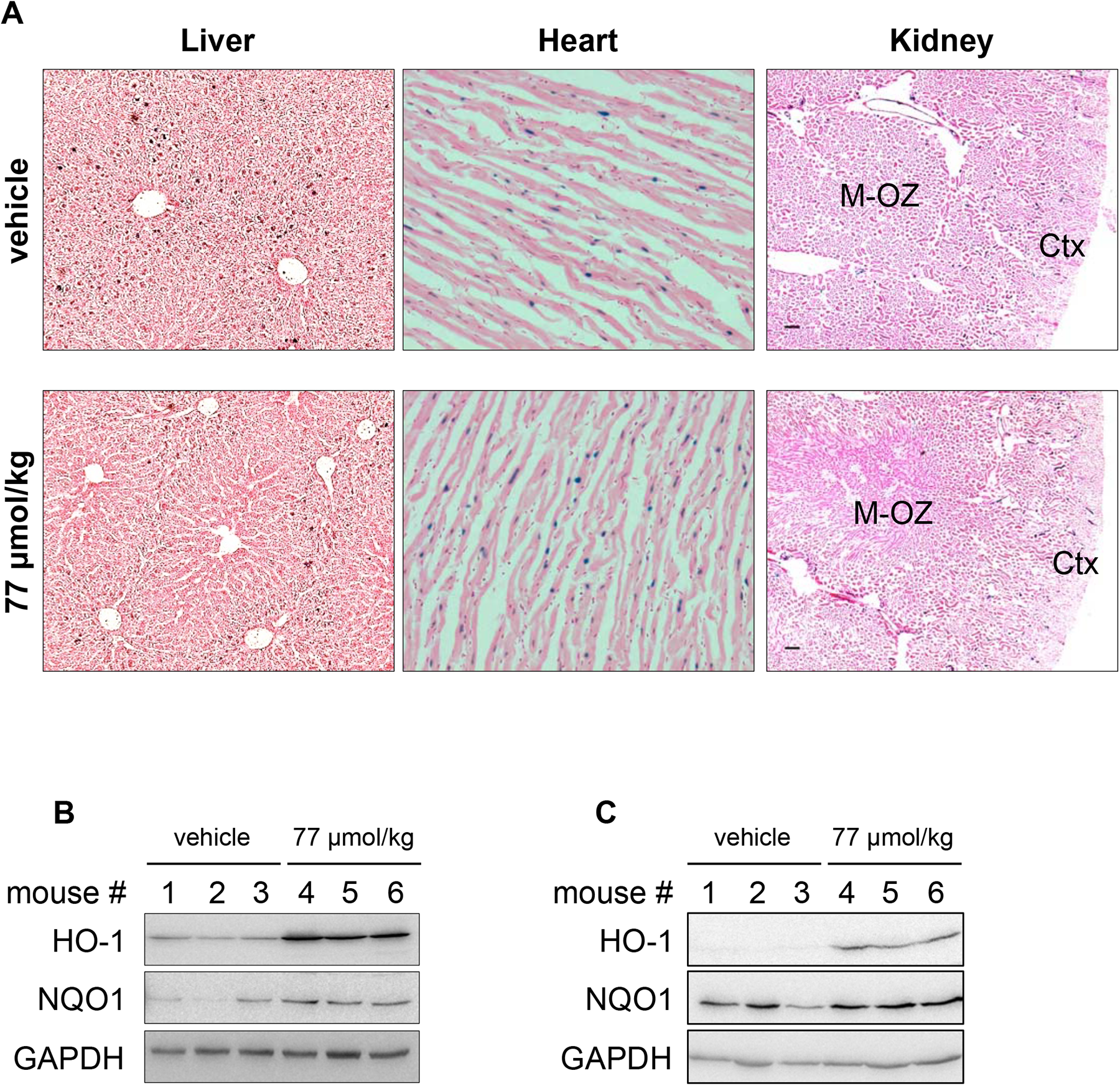
Acute iAs exposure *in vivo* does not increase p21 reporter activity. **A.** Triplicate p21^+/r^ mice (male, 16-18wo) were dosed (p.o.) with 77 μmol/kg iAs. 24h later tissues harvested and β-galactosidase activity assayed as in Figure 1. Representative images are shown for Liver (10x), Heart (20x) and Kidney (2.5x). **B** and **C**. Liver and Kidney lysates were probed for indicated proteins as in Figure 1. Ctx, cortex; M-OZ, medulla outer zone.

Acute dosings are helpful to identify the toxic mechanisms associated to iAs exposure [42]. However, real life exposure is typically chronic from contaminated water or air pollution [48]. To assess the ability of reporter mice to sense stress-induced changes following chronic administration, adult HOTT mice were treated with environmentally relevant concentrations of 50 ppb or 5 ppm iAs in drinking water for 30 days [41]. As a positive control, one HOTT reporter group was exposed to a single dose of 77 μmol/kg (Figure 3 and Figure S3). Interestingly, in HOTT mice exposed chronically to 5 ppm iAs, a positive but sparse induction of the reporter activity was observed in cardiac cells and a robust activation in the kidney medulla (outer zone) comparable to that observed on acute exposure (Figure 3A and B), but not in liver (Figure 3C). The prolonged iAs treatment did not result in overall pathological changes in the liver, as revealed by Haematoxylin/Eosin and Masson’s analysis (Figure S3C and D). Western-blot analysis in kidneys confirmed the increased expression of the HO-1 protein in chronically exposed mice (Figure 3D). NQO1 protein levels remained unchanged. The above results collectively demonstrate the capacity of the HOTT reporter to detect iAs effects on both chronic and acute exposure with the most prominent effects being in kidney, liver and heart tissues.

**Figure 3.**
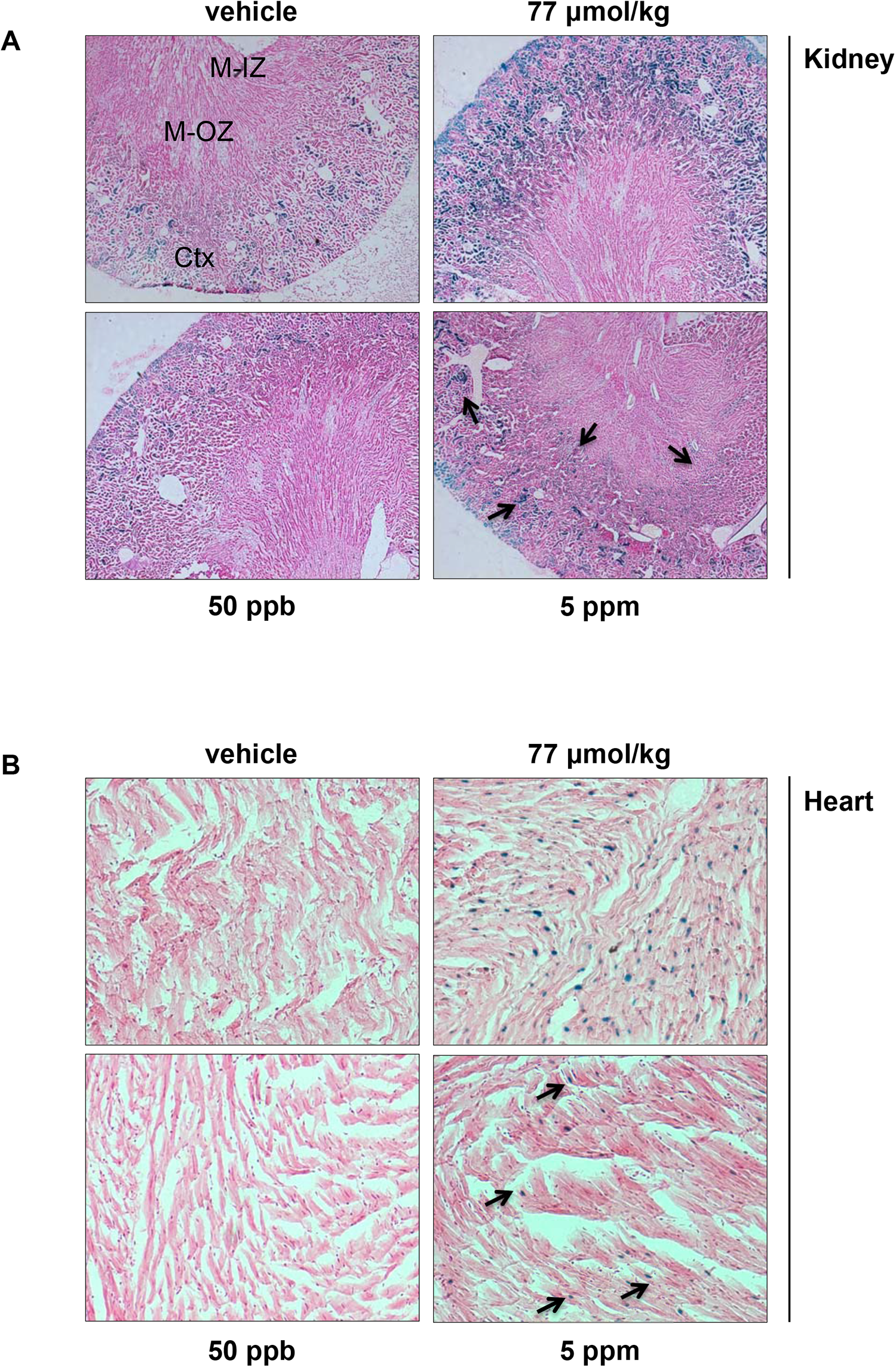
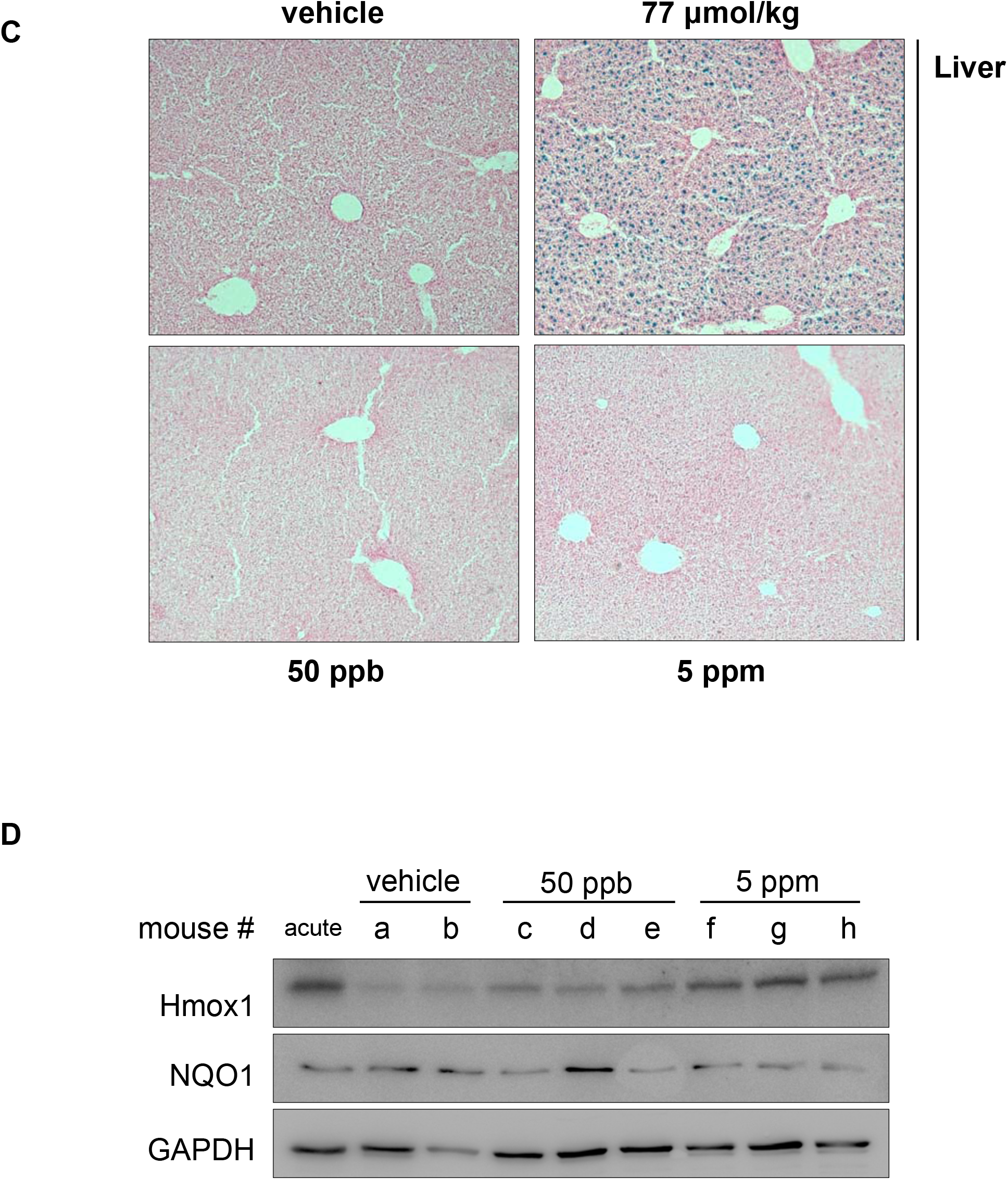
Chronic iAs exposure induces HOTT reporter activation *in vivo*. Four HOTT^+/r^ mice (female, age-matched 16wo) were exposed for 30 days to indicated concentrations of iAs in drinking water (50 ppb or 5 ppm). At end of study, an additional group of aged matched female mice were treated with an acute dose of iAs (77 μmol/kg) and tissues processed as in Figure 1. Representative sections of kidney (**A**, 2.5x), heart (**B**, 20x) and liver (**C**, 10x). Ctx, cortex; M-OZ, Medulla outer zone; M-IZ, Medulla inner zone. **D**, Kidney lysates were probed as in Figure 1. Black arrows indicate β-galactosidase staining.

### Epigenetic analysis of iAs exposed tissues in HOTT mice

In order to establish whether our treatments in mice reflect the molecular events observed in exposed humans, we analysed epigenetic changes associated with iAs exposure in HOTT mice. Changes in the expression and epigenetic regulation of genes associated to different stress responses (e.g. *NFKB1a, SOCS3* or *EGR1)* have been proposed as biomarkers of iAs exposure at the population level [47, 49]. Similar responses have been observed in *in vitro* and *in vivo* laboratory models [50]. We studied the DNA methylation levels of SOCS3, EGR1, JUNB and DUSP1 genes in tissues derived from our cohort of chronically exposed mice. DNA isolated from the heart and liver tissues were subjected to bisulfite conversion and pyrosequencing analysis. Total methylation levels in two different DNA regions and specific CpG positions were determined in these genes (Table 1 and Figure S4B). When compared to unexposed mice, we found a significant hypomethylation in the total DNA methylation of these genes at the second region of study. Interestingly, these changes were observed at the lower iAs concentration and in a tissue specific manner: DUSP1, EGR1 and SOCS3 hypomethylation was observed in heart, but changes in JUNB were observed in liver (Table 1 and Figure S4B).

**Table 1.** Level of total DNA methylation of various genes in arsenic-treated mice. Samples were derived from animals described in Figure 3 and DNA methylation levels quantified as indicated in methods. *,**,***: statistically significant difference from control at p < 0.05, 0.01 and 0.001, respectively.

| Genes | Treatment | Level of DNA methylation (Mean ± SE) | | | |
| --- | --- | --- | --- | --- | --- |
|  |  | 1<sup>st</sup> Study region |  | 2<sup>nd</sup> Study region |  |
|  |  | Heart | Liver | Heart | Liver |
| **DUSP1** | Control | 16.46 ± 0.65 | 7.22 ± 1.74 | 21.35 ± 1.28 | 23.81 ± 1.81 |
| | 50 ppb- 1month | **18.45 ± 0.61\*** | 9.51 ± 1.65 | **11.83 ± 1.18\*\*\*** | 24.07 ± 1.66 |
| | 5000 ppb- 1month | 15.03 ± 1.5 | 7.08 ± 1.21 | **16.80 ± 1.25\*\*** | 18.84 ± 3.01 |
| **EGR1** | Control | 25.02 ± 0.8 | 15.67 ± 4.74 | 15.41 ± 0.72 | 13.07 ± 0.90 |
| | 50 ppb-1month | 26.2 ± 1.26 | 16.63 ± 1.25 | 14.55 ± 0.42 | 12.96 ± 0.59 |
| | 5000 ppb-1month | 24.75 ± 1.42 | 18.71 ± 1.42 | **10.21 ± 0.42\*\*\*** | 12.42 ± 0.37 |
| **JUNB** | Control | 11.01 ± 1.6 | 6.43 ± 1.61 | 21.71 ± 2.76 | 34.01 ± 3.72 |
| | 50 ppb- 1month | 16.30 ± 2.21 | 8.98 ± 1.03 | 17.62 ± 0.61 | **26.77 ± 2.67\*\*** |
| | 5000 ppb- 1month | 14.49 ± 3.44 | 4.27 ± 0.96 | 16.85 ± 2.18 | **19.31 ± 3.08\*\*** |
| **SOCS3** | Control | 30.14 ± 3.51 | 19.85 ± 1.56 | 2.9 ± 0.14 | 14.03 ± 2.78 |
| | 50 ppb- 1month | 27.63 ± 4.79 | 16.24 ± 3.35 | 3.04 ± 0.39 | 8.62 ± 2.88 |
| | 5000 ppb- 1month | 29.68 ± 2.90 | 17.30 ± 3.93 | **2.07 ± 0.08\*\*** | **3.62 ± 1.06\*** |

### NRF2 mediates Hmox1 expression upon *in vivo* iAs exposure

The expression of Hmox1 is controlled by a number of environmentally-regulated transcriptions factors (reviewed in [51]). We have previously show that these different modes of activation can be measured using the HOTT reporter [34]. In order to establish the role of NRF2 (activated by oxidative stress) in the activation of the HOTT reporter upon iAs exposure we crossed the reporter mice onto a NRF2-KO (NRF2 null) background. LacZ staining of untreated NRF2-KO_HOTT animals (12 males, 7 females, aged from 12 to 30 weeks old) was performed. Interestingly, the deletion of NRF2 resulted in a marked variability of reporter basal expression in different tissues and within same individual mice, regardless of age or sex (Figure 4A). In tissues such as brain (hippocampal formation, cerebellum), lung (bronchioles) and kidney (tubules), we observed a diversity of staining, ranging from loss of reporter activity to no difference compared to littermates NRF2wt_HOTT reporter mice. Further investigations will identify the source of this variability.

**Figure 4.**
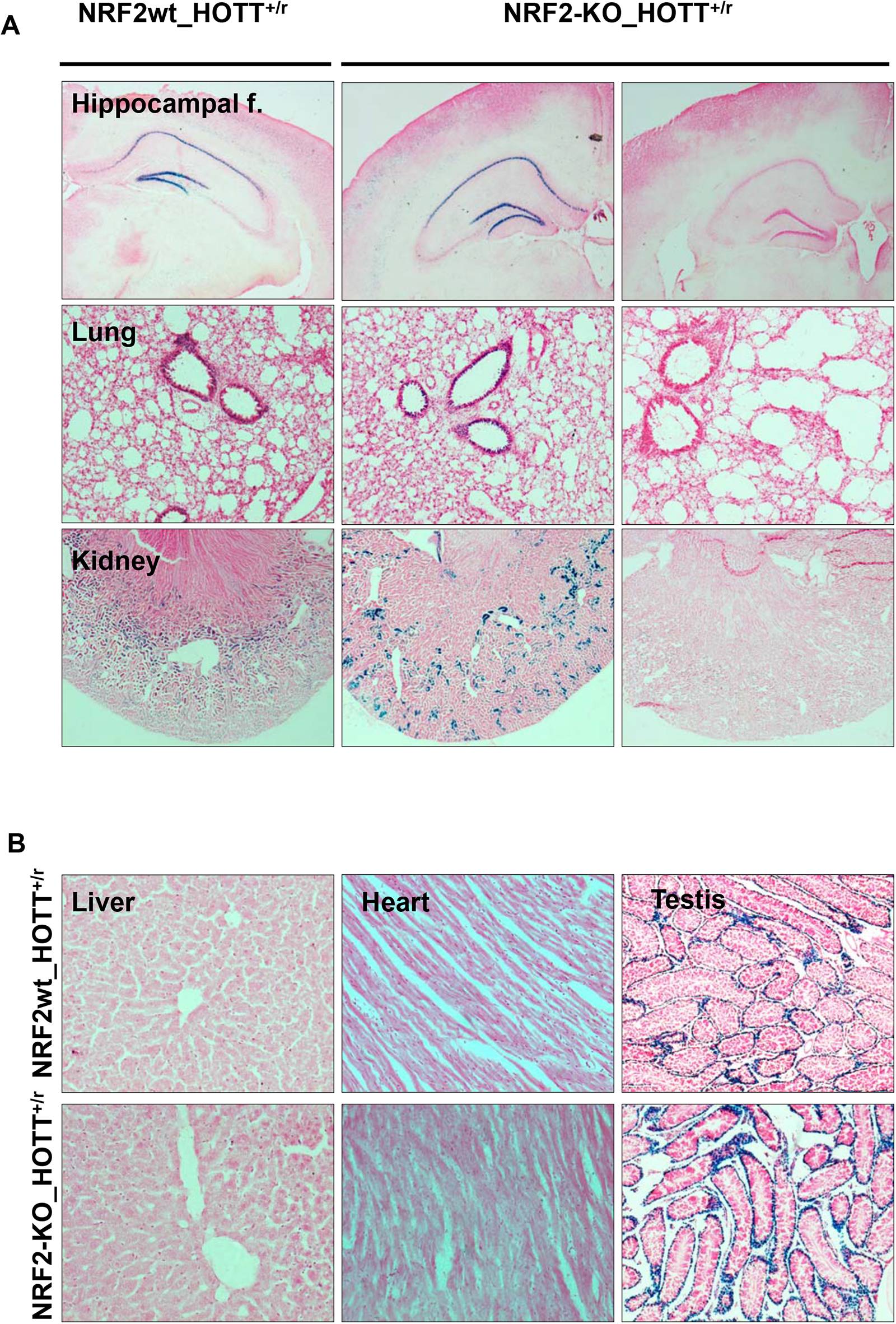
Basal Hmox1 expression in a novel NRF2-KO_HOTT reporter model. **A, B.** Indicated tissues were harvested and *in situ* β-galactosidase assay performed from untreated animals. Representative images are shown for hippocampal formation (2.5x), lungs (10x), Kidney (2.5x), heart (20x), liver (10x) and testis (5x). n = 12 males, 7 females, 15 ± 3 weeks old.

As NRF2 regulates the cellular redox homeostasis, we considered the possibility that in NRF2-KO mice, there might be a general decrease in antioxidants. Indeed, electron paramagnetic resonance (EPR) imaging studies have demonstrated decreased tissue-reducing activity in NRF2-KO in comparison with wild-type mice, which is particularly evident in aged animals [52]. This could be reflected in the induction of the HO-1 reporter activity. The absence of NRF2 did not induce HOTT reporter activity in tissues where the basal reporter activity was not detected previously, such as liver or heart (Figure 4B) and large intestine (Figure 5B). Furthermore, the expression of the HOTT reporter in testis was not altered by the absence of NRF2. These results suggest that either the absence of NRF2 alone is insufficient to cause oxidative stress, or that NRF2 is required for the expression of the HOTT reporter. The normal phenotype and lifespan of the NRF2-KO mouse under standard laboratory conditions (i.e. in the absence of any challenges) supports the second possibility.

**Figure 5.**
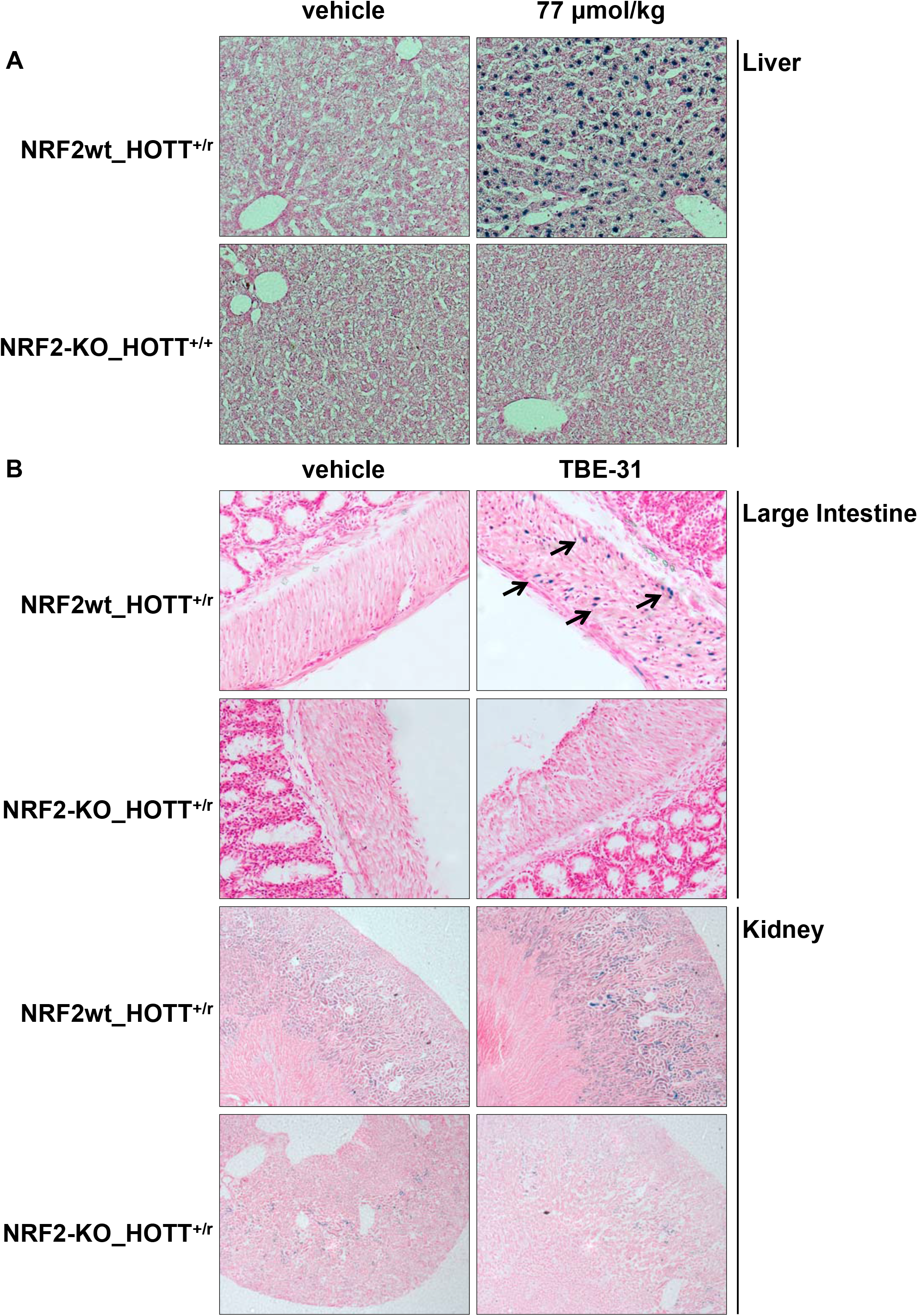
Characterization of a novel NRF2-KO_HOTT reporter model. **A.** Triplicate NRF2wt_HOTT^+/r^ or NRF2-KO_HOTT^+/+^ mice were treated with vehicle or 77 μmol/kg iAs. 24h later tissues were harvested and *in situ* β-galactosidase assay performed in liver (20x). **B.** Triplicate mice NRF2wt_ HOTT^+/r^ or NRF2-KO_HOTT^+/r^ were treated with vehicle or TBE-31 as described in methods. 24h later tissues were harvested and *in situ* β-galactosidase assay performed in the indicated tissues. Representative images are shown for large intestine (20x) and kidney (2.5x).

The NRF2-KO mice were generated by replacing exon 5 of the *Nfe2l2* gene, which encodes the DNA binding and dimerization domains, with an NLS-LacZ-neo (nuclear localization signal-LacZ-neomycin resistant gene) cassette [37, 53]. To ensure that any LacZ activity detected in our studies is derived from the HOTT reporter and not from the targeted *Nfe2l2* alleles, we exposed NRF2-KO_HOTT^+/+^ and Nrf2wt-HOTT^+/r^ mice to an acute dose of iAs (Figure 5A). Neither control nor iAs exposed liver or kidney tissue derived from NRF2-KO_Hmox1^+/+^ mice had β-galactosidase activity, confirming that the NLS-LacZ-Neo cassette does not contribute to the signal detected in subsequent experiments. To demonstrate the utility of our NRF2-KO_HOTT reporter in detecting specific NRF2-dependent Hmox1 transcriptional activation, we treated NRF2wt_HOTT^+/r^ or NRF2-KO_HOTT^+/r^ mice with vehicle or TBE-31, a potent NRF2 activator. TBE-31 treatment activated the HOTT reporter in the kidney cortex and smooth muscle cells of the large intestine (Figure 5B). This reporter activation was dependent on NRF2, as it was absent in NRF2-KO_HOTT^+/r^ mice. Therefore, the combination of NRF2wt_HOTT^+/r^ and NRF2-KO_HOTT^+/r^ mice provides a novel approach to define the role of NRF2 in the transcriptional regulation of Hmox1.

To investigate the role of NRF2 in the context of HOTT reporter activation by iAs, NRF2-KO_HOTT^+/r^ or NRF2wt_HOTT^+/r^ mice were exposed to an acute dose of iAs (Figure 6). In NRF2-KO mice the reporter activity was markedly attenuated in the kidneys, heart and liver. This indicates that NRF2 is the main transcription factor activated by arsenic *in vivo* which results in the induction of Hmox1. No HOTT reporter activation was observed in any other tissues examined, indicating that NRF2 deletion does not promote the activation of other stress pathways which regulate HO-1 by iAs exposure.

**Figure 6.**
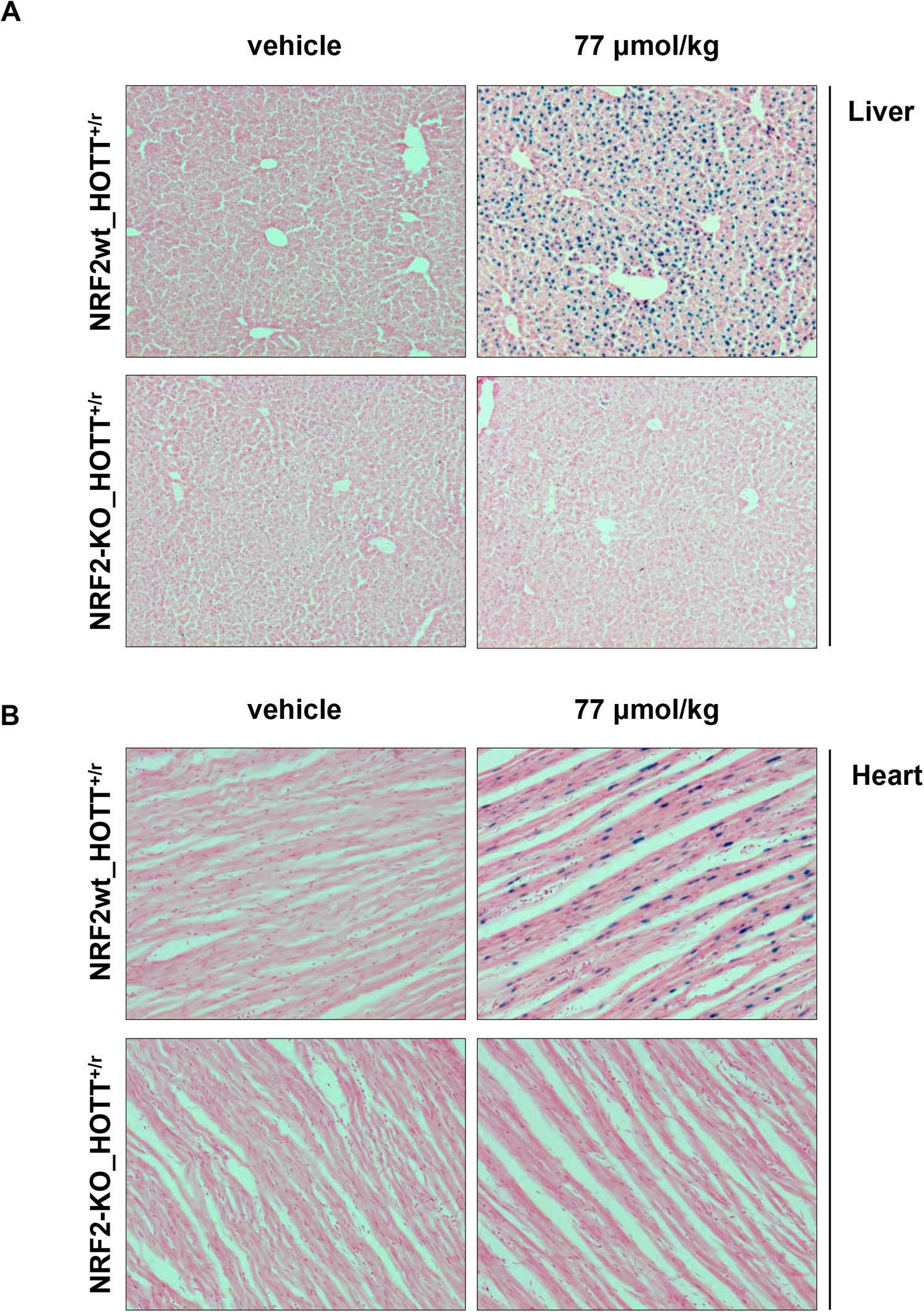
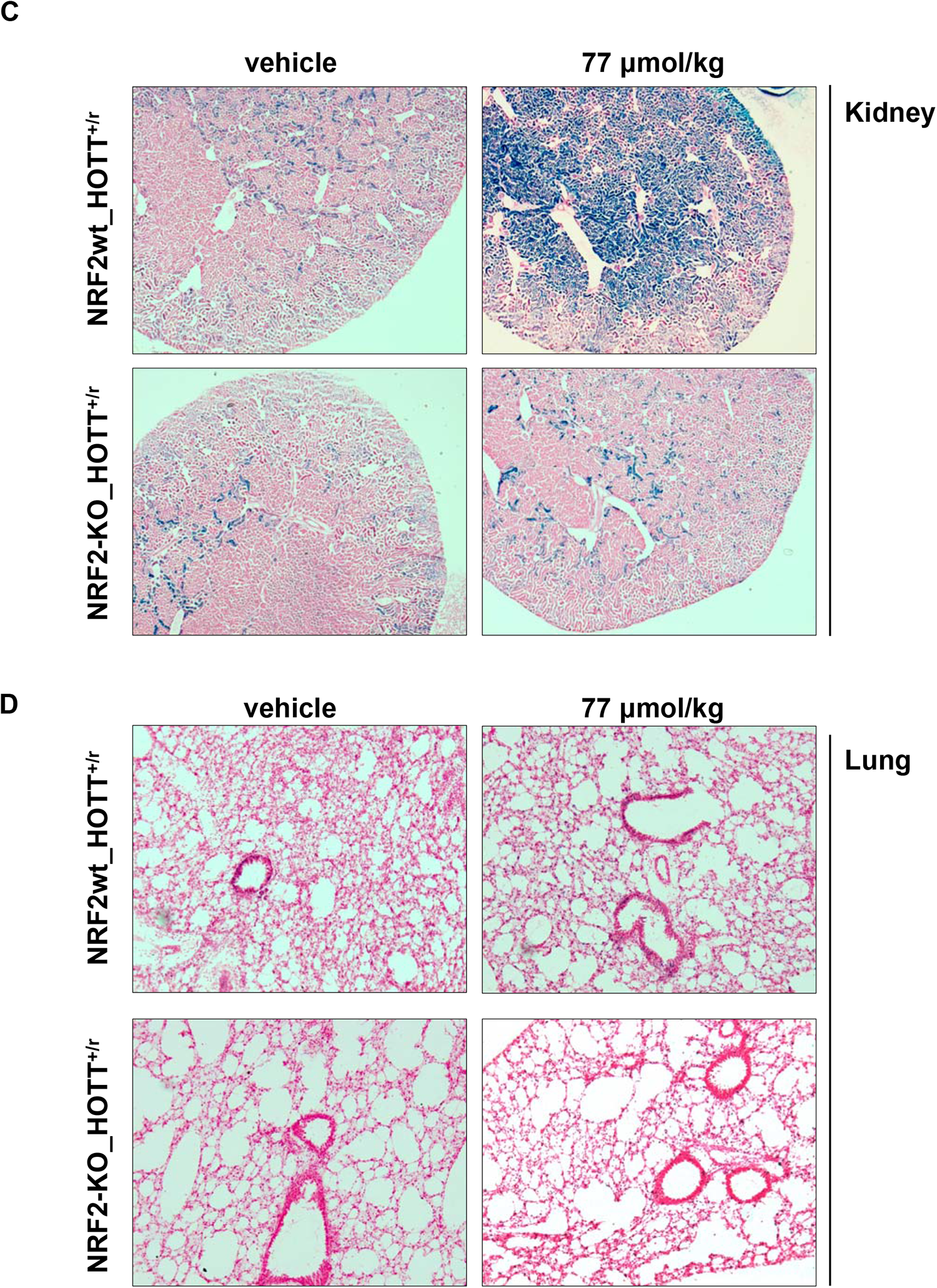
NRF2 mediates the HOTT reporter activation in response to acute iAs exposure. Triplicate NRF2wt_HOTT^+/r^ or NRF2-KO_HOTT^+/r^ mice were treated with vehicle or 77 μmol/kg iAs. 24h later tissues were harvested and *in situ* β-galactosidase assay performed in liver (**A**, 10x); heart (**B**, 20x); kidney (**C**, 2.5x); or D, lungs (**D**, 10x).

### Utility of the HOTT reporter to identify modulators of the toxic effects of arsenic

In light of the above results and to test the utility of the HOTT reporter as a surrogate biomarker of iAs toxicity *in vivo*, mice were treated with an acute dose of iAs in combination with the antioxidant *N*-acetyl cysteine (NAC). 24h after dosing, and before tissue harvesting, we examined the reporter activation by whole-body imaging (Figure S5B). No clear luminescence pattern was observed in animals treated with arsenic. As expected, β-galactosidase activity analysis demonstrated induction of reporter activity in heart, liver and kidney. This activity was markedly reduced in all tissues examined on the coadmistration of NAC (Figure 7A-C), indicating protection from the iAs-induced oxidative stress. Quantitation of the β-galactosidase signal by densitometry indicated a significant level of protection in all tissues examined (Figure 7D). These data indicate that the HOTT reporter could provide a valuable *in vivo* model to identify treatments which reduce arsenic toxicity.

**Figure 7.**
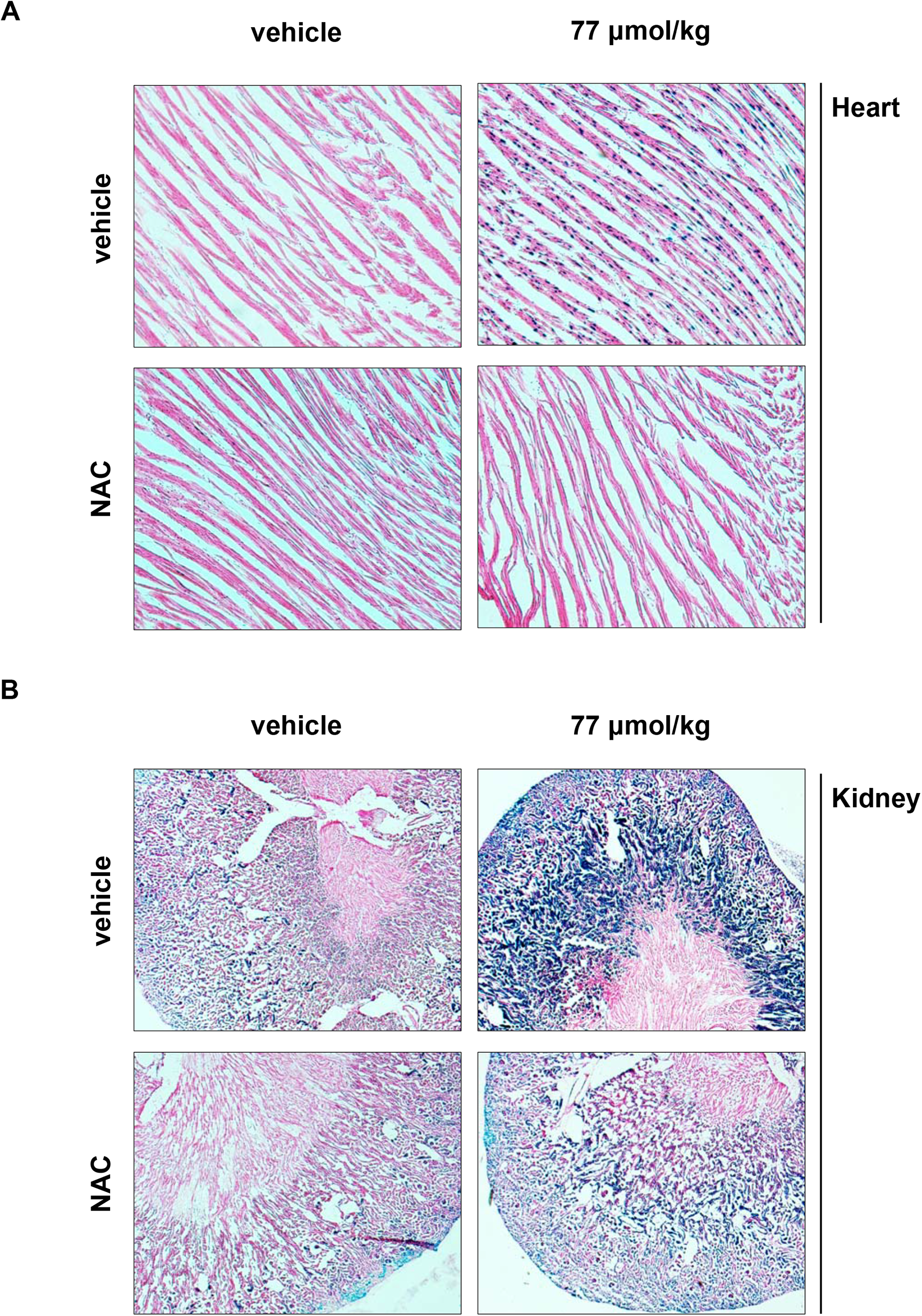
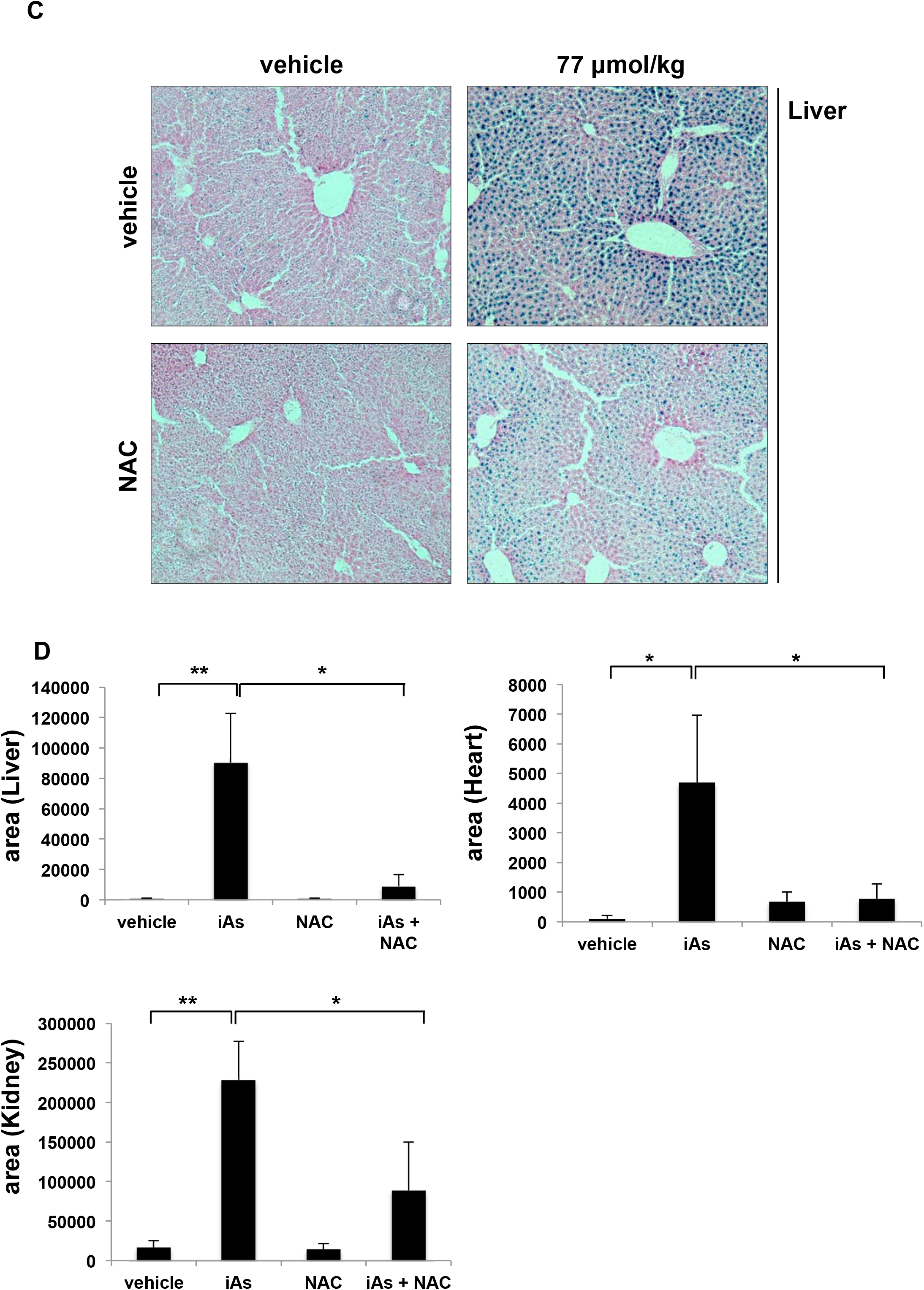
HOTT reporter predicts therapeutic utility on iAs toxicity. Triplicate HOTT^+/r^ mice were treated with vehicle, 77 μmol/kg iAs, 300mg/kg NAC or a combination of iAs and NAC. 24h later tissues were harvested and *in situ* β-galactosidase assay performed in heart (**A**, 10x); Kidney (**B**, 10x); Liver (**C**, 10x). **D**, densitometry evaluation (Figure 8A-C) of microscopic images. Data represented as mean + SD. T-test performed, * p<0.05; ** p<0.01.

## Discussion

In this paper we describe the application of three *in vivo* reporter models p21, HOTT and NRF2_KO-HOTT to detect the cellular stress induced by the environmental toxin iAs. Interestingly, iAs markedly induced the HO-1 reporter, but did not activate the p21 DNA damage/p53 reporter. We used a combination of genetic and pharmacological approaches to demonstrate *in vivo* that oxidative stress is the initiating cause of arsenic poisoning in a tissue specific manner [54]. The absence of murine p21 induction is interesting in the light of the evidence that iAs exposure has the capacity to induce DNA damage in exposed infants [44]. Reporter activation occurred in the absence of overt liver tissue damage as measured by H&E, Masson’s and biochemical analysis so provides an early biomarker of the iAs toxicity. Importantly, additionally to a NRF2 response, we observed specific changes in the DNA methylation levels of genes related to the regulation of the immune system or cellular proliferation [9, 49] and in a tissue-specific manner. These findings support the utility of mice studies to understand biochemically the iAs associated toxicity and to inform future preclinical studies.

The capacity of iAs to induce stress responses was studied using four iAs dosing regimens; acute and chronic exposures at low and high doses. A comprehensive analysis of different organs revealed a tissue-specific HOTT reporter activation, occurring mainly in liver, kidney and heart. The oxidative stress response measured with the HOTT reporter reflects the previously observed dynamic, dose-dependent accumulation, metabolism and distribution of arsenicals in different organs [27, 55]. In mice and humans, after absorption in the gastrointestinal tract, iAs is widely distributed before being excreted in the urine in a biphasic pattern. The majority of iAs is eliminated in the first 4h (t_1/2α_ = 2h) with rest being eliminated within 4 days (t_1/2β_ = 44.6h) [56]. The heart (i.e. cardiomyocytes) was particularly sensitive to iAs, as reporter activation was detected after acute exposure to 5.4 μmol/kg and 77 μmol/kg and on chronic exposure to 5 ppm. It has been reported that on acute iAs exposure, there is a significant accumulation in the heart, which was double the concentration in the liver and comparable to that in the lung, spleen or skin. Moreover, the same exposure in knock out mice for aquaglyceroporin-9 (a major transporter of iAs species), evidenced that heart accumulated 10-20 times more arsenic than wild type, the highest rate of iAs accumulation for any measured tissue [57].

In the kidneys, HOTT reporter activation was detected at a high acute iAS dose and on chronic treatment (5 ppm). Interestingly, no reporter activation was observed in the kidneys of mice treated with an equivalent acute dose of 5.4 μmol/kg. This raises the possibility of the induction at 5 ppm is due to an accumulation of iAs species. This hypothesis is supported by reports on chronic iAs exposure and steady-state (typically after nine repeated daily doses) tissue accumulations studies, where it has been described that kidneys accumulate the highest levels of iAs species [56]. Data on the heart was not reported in these studies. In liver, HO-1 induction was only detected in mice exposed to 77 μmol/kg. iAs is rapidly converted in the liver to mono methyl arsenic (MMA) and dimethyl arsenic (DMAs; the main excreted form) in a reaction catalyzed by As(+3) methyltransferase. DMAs are then accumulated in different tissues and may contribute to the iAs toxicity [58]. Further research will be required to establish why arsenic exposure in other tissues, such as lung, large intestine, spleen and brain, did not activate the reporter.

The HO-1 promoter can be activated by a number of transcription factors including those involved in inflammation [51]. Using the NRF2-KO_HOTT model we demonstrated that the induction of HO-1 by iAs was mediated by NRF2. This transcription factor is mainly activated by toxic electrophiles and oxidative stress [59], providing evidence that the induction of oxidative stress is the mechanism of *in vivo* HO-1 activation. This is further substantiated by the finding that administration of the antioxidant NAC significantly reduced the expression of the reporter in all the affected tissues. A role for NRF2 in HO-1 activation following exposure to iAs in vivo has been previously suggested [60]. Different mechanisms have been proposed to explain the activation of NRF2 by iAs, both dependent and independent of Keap-1 oxidation [61, 62]. Following a single iAs dose, NRF2 stabilization and transcriptional activity *in vivo* peaks at 12h, decreasing to basal levels at 24-36h [43, 63]. This is reflected in the Hmox1 mRNA levels, which follow a similar pattern of expression, peaking at 4h and returning to basal levels after 12h. These findings highlight one of the advantages of our reporter system as the half life of β-gal is ~36h [64], which allows transcriptional activation to me measured even if very few cells are affected.

Different reports suggest a protective role for NRF2 against iAs toxicity. For example, NRF2-KO mice exposed for 4 months to 5ppm are more vulnerable to bone loss than their wild type counterparts [65]. Similarly, exposure of NRF2-KO mice for six weeks to 10ppm or 100ppm arsenic concentrations resulted in a more severe pathological changes in the liver and bladder, compared to wild type mice [31]. Consistent with these findings, we found that exposure to 5 ppm iAs for 4 weeks activated of the HOTT reporter in heart and kidney, although to a lesser extent to acute exposures. This result suggests that sustained oxidative damage in these critical tissues occurs. Chronic activation of NRF2 due to prolonged oxidative stress has been linked to many pathologies, including cardiovascular defects, cancer and chemo-resistance. It is tempting to speculate that the observed chronic oxidative stress (in *in vivo* studies and at the population level) could be contributing to the pathologies found in iAs exposed populations [54], suggesting that activation of the NRF2 system may provide a means of reducing iAs toxicity.

## Conclusion

We demonstrate that stress reporters provide a robust and highly reproducible approach to study the deleterious processes induced by iAs and potentially other compounds or compound mixtures. The reporter approach has a number of advantages over other currently used approaches. For example, β-galactosidase activity (LacZ staining) allows the detection of oxidative stress in conditions of chronic exposure to low iAs concentrations, at single cell level. This is not possible using the previously described NRF2 reporter mice which were only able to detect low resolution changes in bioluminescence at high iAs concentrations [66]. Other methods for measuring oxidative stress, such as changes in 8-OHdG or blood GSH levels, are transient and only detect oxidative damage at high levels of exposure and are therefore of limited utility. Relative to conventional toxicological studies this model allowed us to use a smaller number of mice in each experimental group (typically, n = 3 vs. n ≥ 5) [27, 42]. Moreover, the use of different stress reporter models allows mechanistic insights into pathways of chemical toxicity to be obtained in a tissue and cell specific manner. The Hmox-1 reporter therefore provides a powerful approach to identify compounds capable of preventing arsenicosis and allows PK/PD relationships for such compounds to be obtained and used to design clinical trials.

## Additional Information

### Author contributions

F.I.V.: funding acquisition, conceptualization, data curation, formal analysis, investigation, project administration, methodology, writing original draft, review and editing.

C.J.H. and C.R.W.: funding acquisition, conceptualization, project administration, review and editing.

T.G.F.: methodology.

M.R.; P.N.: DNA methylation investigation, resources, data curation, analysis and methodology.

A. T. D-K.; T.H.: resources, review and editing.

## Acknowledgments

We thank the Wolf/Henderson lab for assistance and discussions, the Medical School Resource Unit staff, Ngaire Dennison (Named Veterinary Surgeon) and University of Dundee Biological Services for technical support and ensuring animal welfare at all times.

---

**Figure S1.**
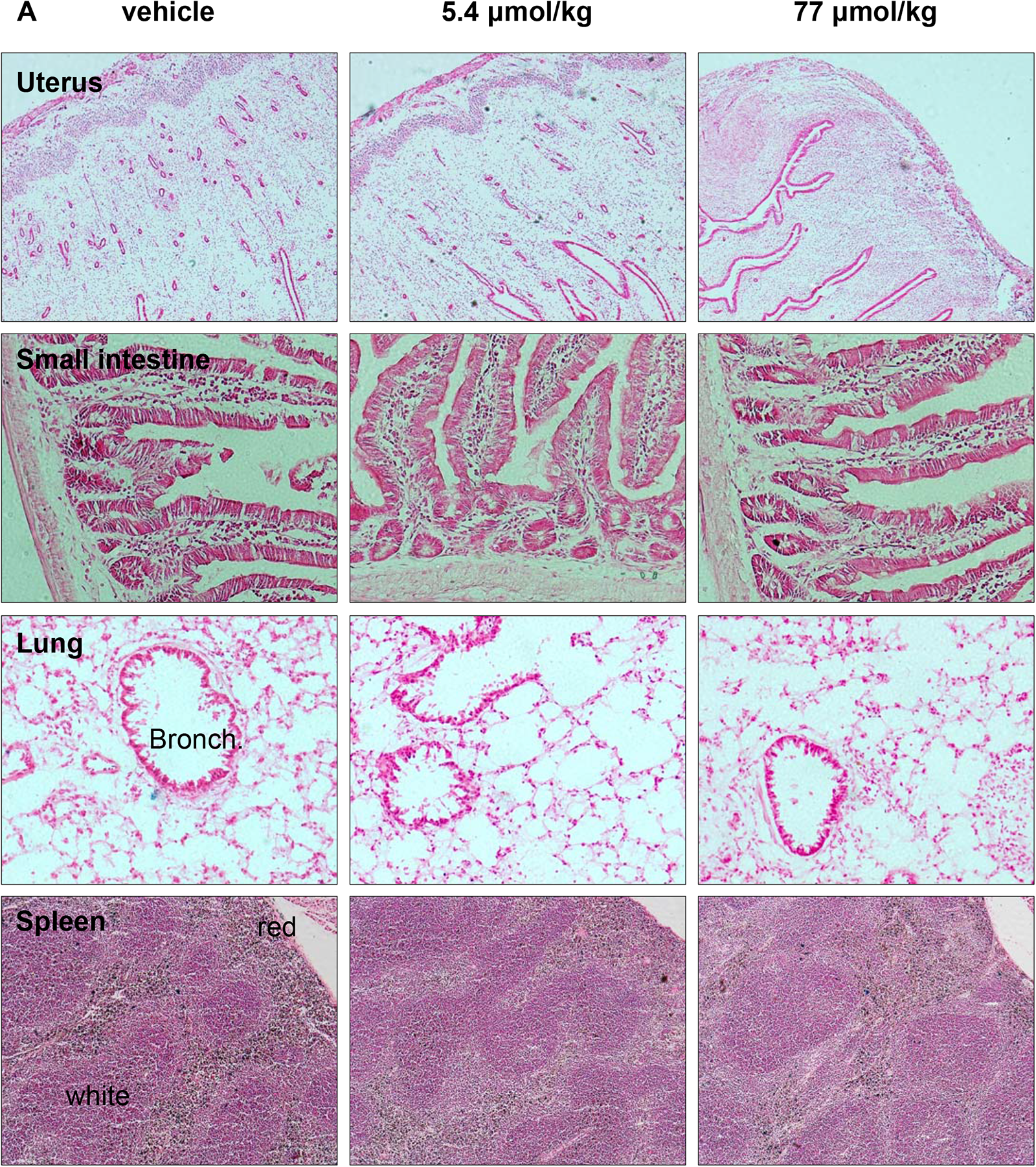
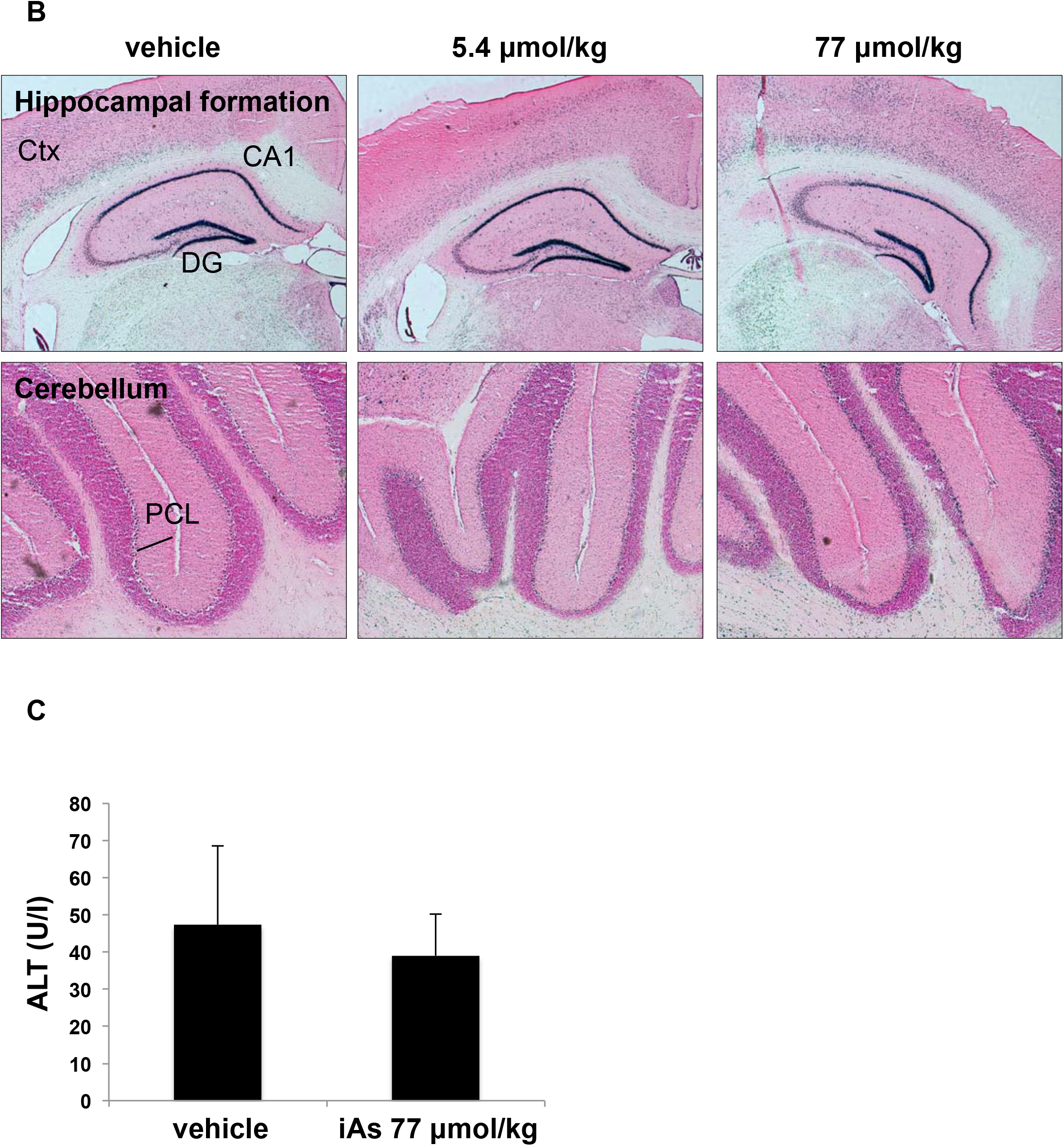
Tissue specific susceptibility to iAs exposure *in vivo* using HOTT reporter mice. **A, B.** Animals and *in situ* β-galactosidase assay performed as in Figure 1, but representative sections of uterus (5x), small intestine (20x), lung (20x), spleen (5x), hippocampal formation (2.5x) and cerebellum (5x) tissues are displayed. Bronch, bronchiole. White, white pulp; Red, red pulp; Ctx, cortex; DG, dentate gyrus; PCL, Purkinje cell layer. **C.** ALT activity in plasma obtained at necropsy (average of 3 mice/treatment).

**Figure S2.**
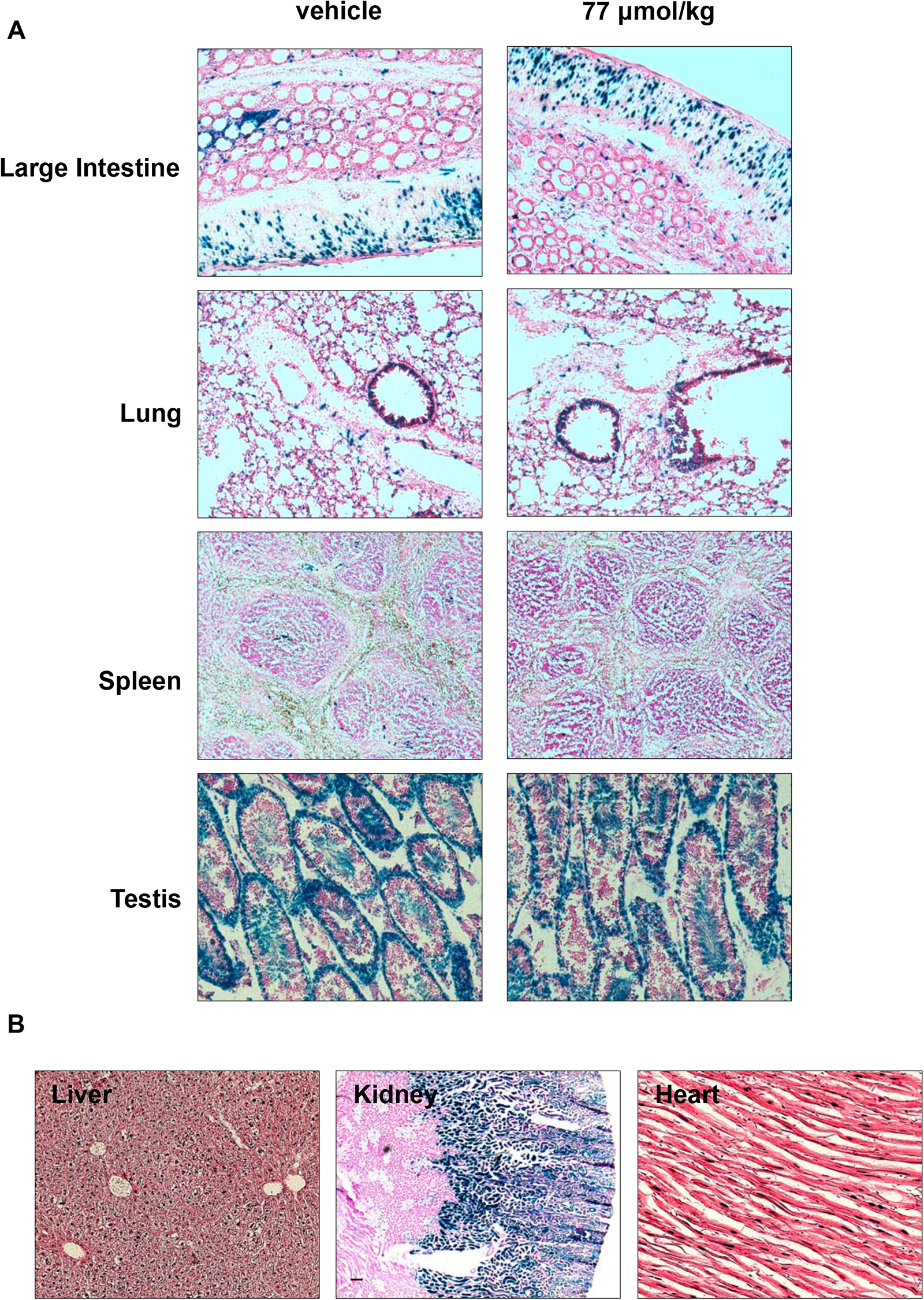
Acute iAs exposure *in vivo* does not increase p21 reporter activity. **A.** Animals and *in situ* β-galactosidase assay performed as in Figure 2, but representative sections of indicated tissues. Representative images are shown for large intestine (10x), lung (10x), spleen (5x) and testis (10x). **B.** Two HOTT^+/r^ male reporter mice where dosed with iAs (77 μmol/kg, p.o.). 24h later tissues harvested and β-galactosidase activity assayed in parallel to animals in Figure 2. Representative images are shown for liver (10x), kidney (10x) and heart (20x).

**Figure S3.**
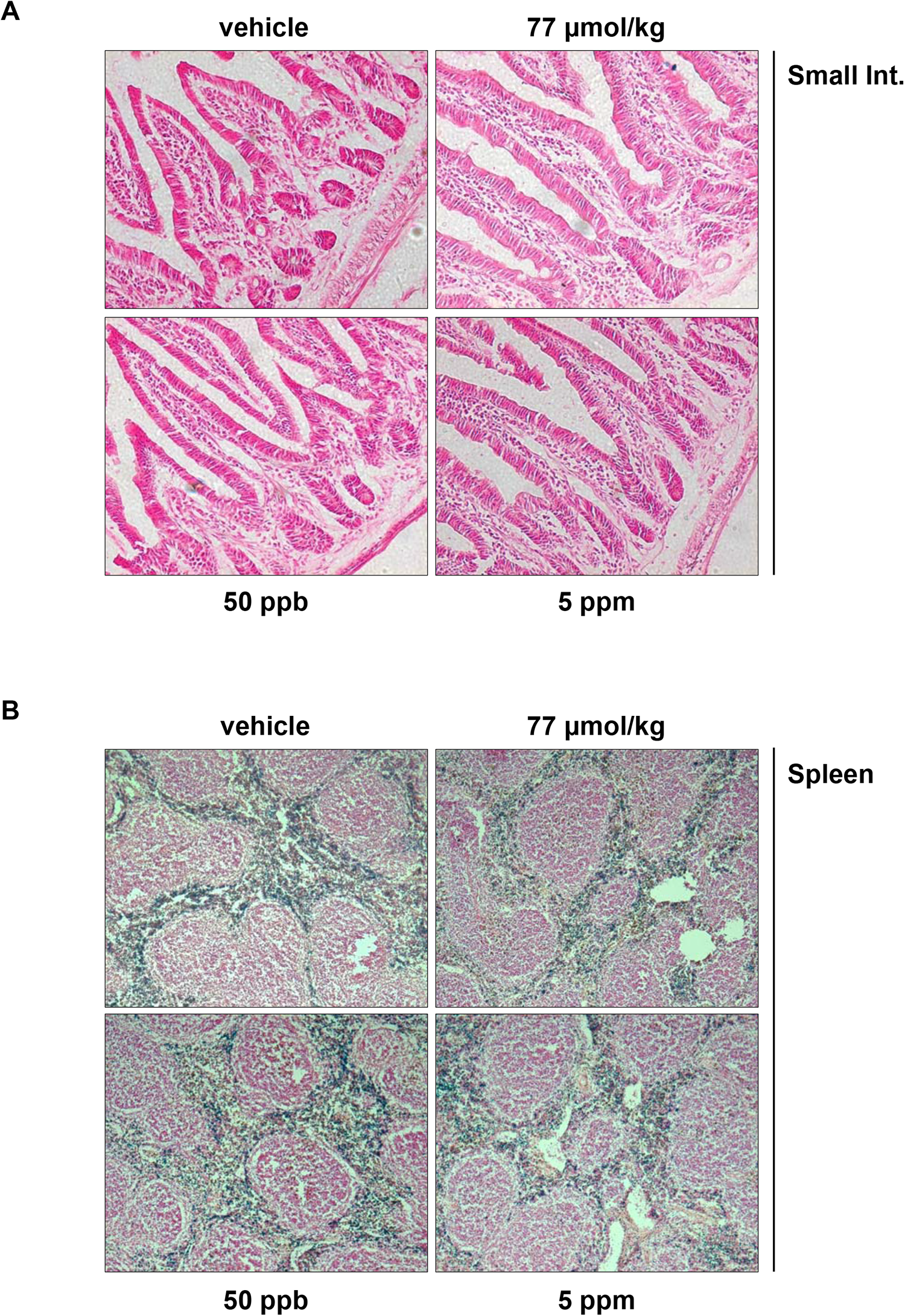
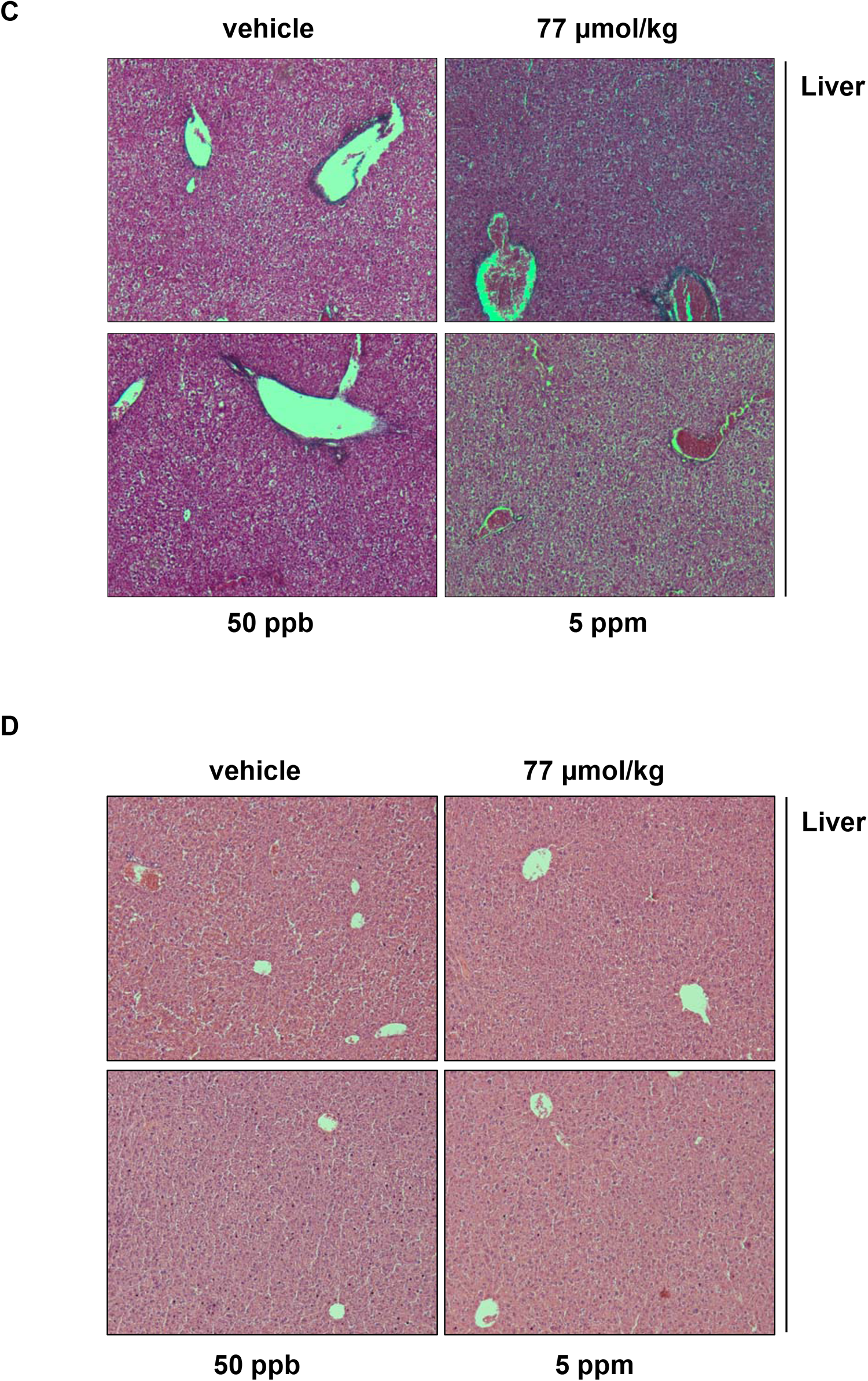
Chronic iAs exposure induces HOTT reporter activation *in vivo*. Animals and *in situ* β-galactosidase assay performed as in Figure 3, but representative sections of indicated tissues: small intestine (**A**, 20x), spleen (**B**, 5x). Liver samples were processed for Masson’s (**C**) and Haematoxylin/Eosin (**D**) staining. Representative images are shown (10x).

**Figure S4.**
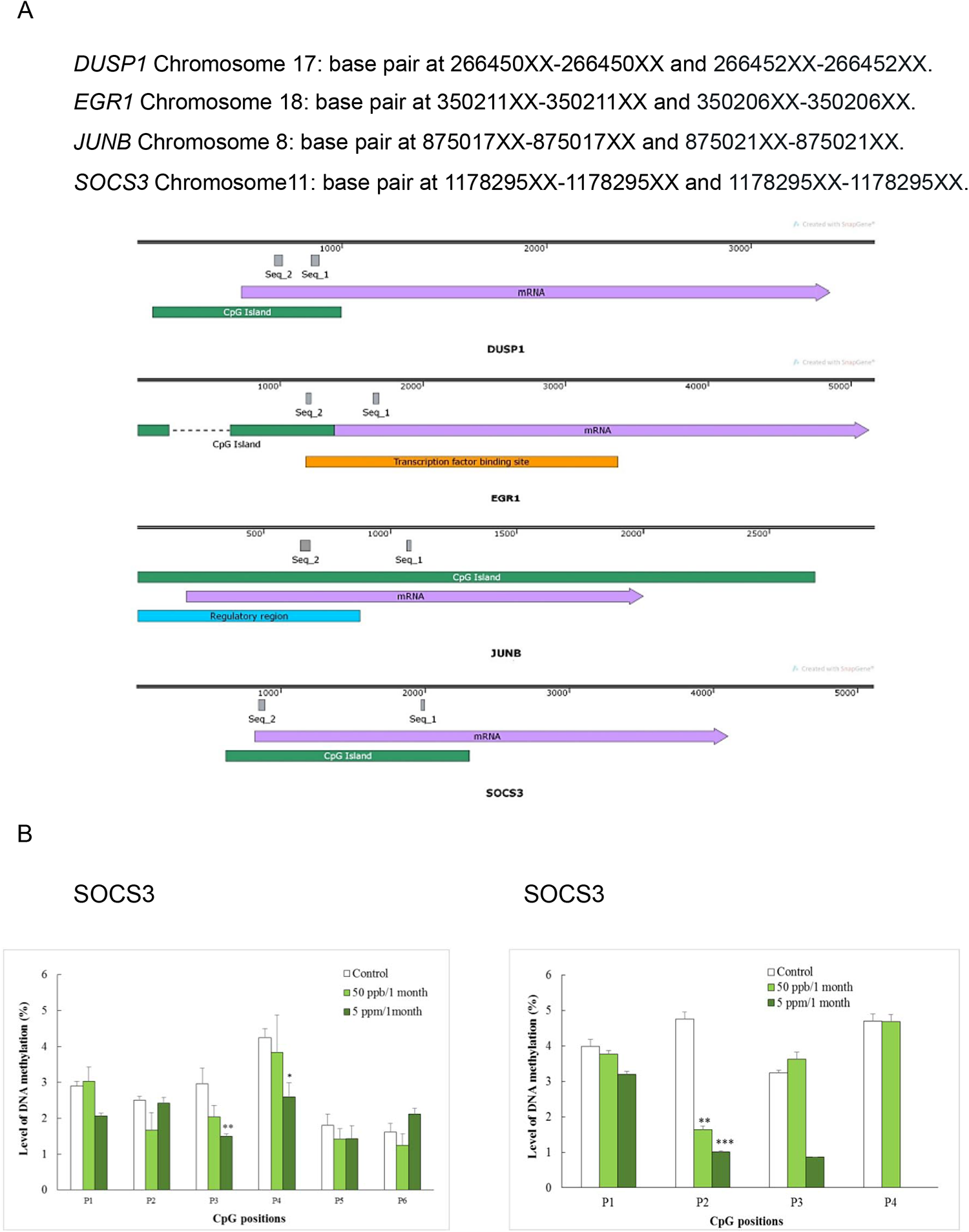
Level of total DNA methylation of various genes in arsenic-treated mice. **A.** The study regions of selected genes using two commercially predesigned CpG assays. **B.** Levels of DNA methylation of *SOCS3* at the second study region (base pair 1178295XX-1178295XX) in heart (left panel) and liver (right panel). Samples were derived from animals described in Figure 3. *,**,*** indicates a statistically significant difference from control at p < 0.05, 0.01 and 0.001, respectively.

**Figure S5.**
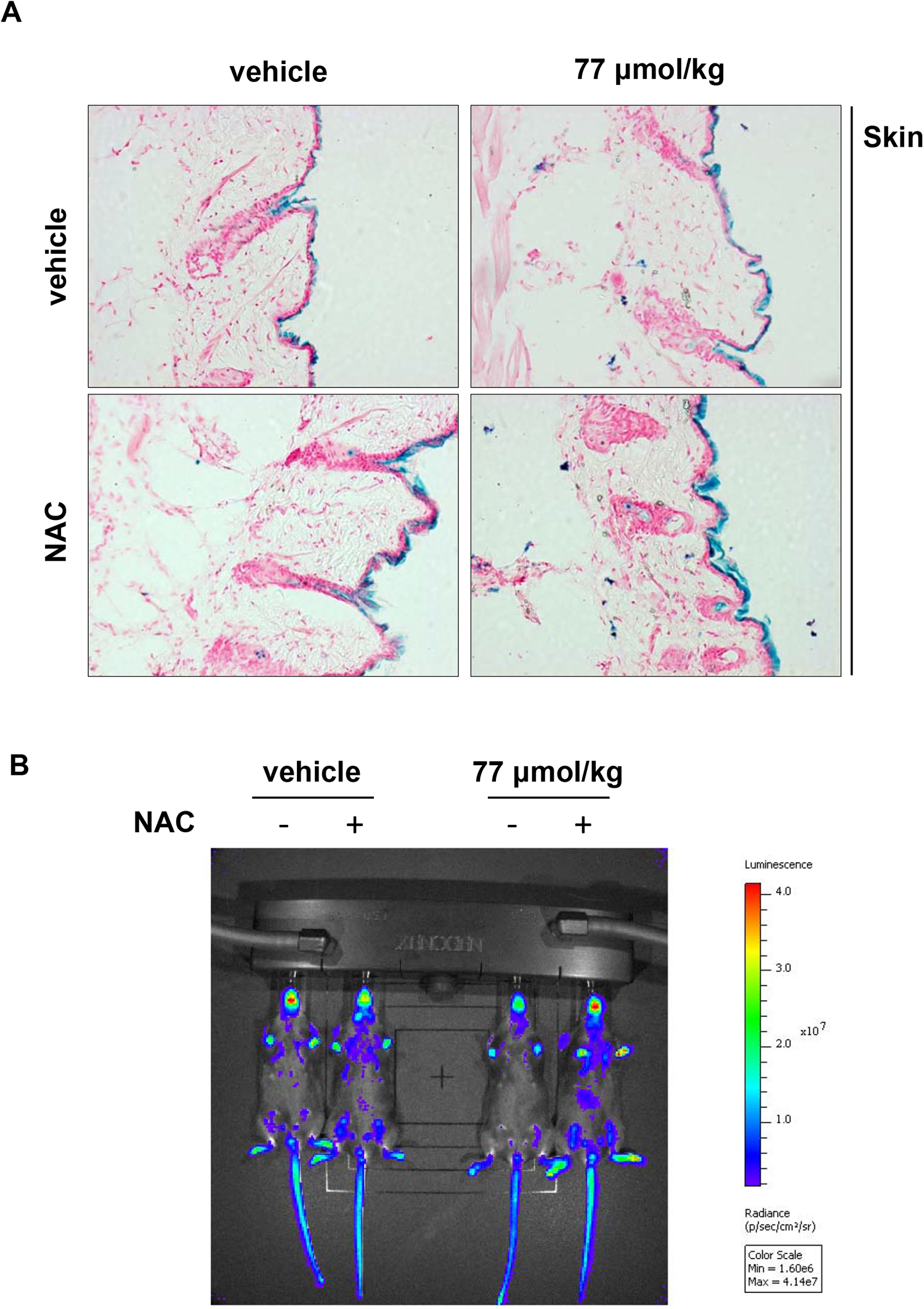
HOTT reporter predicts HOTT reporter predicts therapeutic utility on iAs toxicity. **A.** Animals and *in situ* β-galactosidase assay performed as in Figure 7, but representative sections of skin tissue (20x) are shown. **B**. *In vivo* bioluminescence images of mice after indicated treatments.

